# Viral replication and host response in airway epithelial cells predicts coronavirus pathogenicity and identifies HCoV-229E as an atypical seasonal coronavirus

**DOI:** 10.64898/2026.11.08.765206

**Authors:** Matthew J Gartner, Arada Hirankitti, Monika Mohenska, Stephany Sanchez, Guizhi Sun, Joseph Chen, Kevin Tang, Nikita Deshpande, Randy Suryadinata, Stephen Wilcox, Daniel P Rawlinson, Sarah L. Londrigan, Daniel Steinfort, Philip Robinson, Jose M Polo, Annette Fox, Jessica A Neil, Kanta Subbarao

## Abstract

Human coronaviruses (HCoVs) range from mild seasonal strains to pathogenic viruses that cause severe pneumonia. The virological and host determinants that distinguish pathogenic coronaviruses (MERS-CoV, SARS-CoV-2) from seasonal strains (HCoV-229E, HCoV-NL63 and HCoV-OC43) remain poorly defined. To identify determinants of pathogenicity, we systematically compared replication, cellular tropism, innate responses and cytopathogenicity of pathogenic compared with seasonal HCoVs across nasal, bronchial, small-airway and embryonic stem cell-derived alveolar type II (AT2) epithelial cell cultures. All HCoVs preferentially infected ciliated cells in the upper airway, but only pathogenic viruses replicated to high titres in AT2 cells. Pathogenic HCoVs reprogrammed AT2 function and suppressed key innate immune signatures, consistent with distal-lung epithelial dysfunction. In contrast, seasonal HCoVs were associated with limited alveolar replication and milder disruption of AT2 homeostasis. Strikingly, despite modest replication in all cell types, HCoV-229E induced pronounced epithelial injury, cilia loss and strong interferon-stimulated gene expression, as well as induction of cellular stress/death programmes resembling those induced by MERS-CoV. These findings highlight the striking differences in cellular responses throughout the airways and between distinct HCoV strains and emphasizes the importance of assessing distal lung tropism and epithelial host responses to evaluate emergent coronaviruses to inform pandemic preparedness strategies.

## Introduction

In the past two decades, three human coronaviruses (HCoVs) have emerged from animals to successfully cause human infections associated with significant public health emergencies. SARS-CoV emerged in Guangdong, China in 2002 and spread to 29 countries, infecting 8,098 people with a case fatality rate (CFR) of 9.5% (*1*). MERS-CoV was first detected in Saudi Arabia in 2012 and has since spread to 27 countries, infecting 2,649 people with a CFR of 36.2% (*2*). In 2019, SARS-CoV-2, the agent responsible for coronavirus disease 2019 (COVID-19), emerged in Wuhan, China, resulting in a global pandemic that has led to more than 7 million deaths (*3*). In addition, there are four seasonal HCoVs (HCoV-229E, HCoV-NL63, HCoV-OC43 and HCoV-HKU1) that also likely emerged from animals in the past and have been endemic in the human population for decades, accounting for 8-25% of common cold cases (*4–6*). While seasonal HCoVs are largely associated with self-limiting infections with mild respiratory symptoms, these viruses can lead to more severe respiratory disease in neonates, the elderly and immunocompromised individuals (*7–11*). Understanding the virological and host determinants that drive the divergent clinical outcomes during infection with the different HCoV strains has key implications for clinical management, therapeutic development and pandemic preparedness.

There is considerable evidence implicating host factors such as age, genetics, co-morbidities and pre-existing immunity with HCoV infection susceptibility and disease severity (*12*). However, it is still unclear why some HCoVs show increased pathogenicity (SARS-CoV, MERS-CoV and SARS-CoV-2) while others appear to have remained relatively mild (seasonal HCoVs). The long period of endemicity of seasonal HCoVs means there is a high degree of population-level immunity to these viruses that likely contributes to amelioration of disease severity (*13, 14*). Factors such as tissue and cellular tropism, host innate and adaptive immune responses and viral antagonism of host cell responses likely also play a role in the pathogenic potential of these viruses. Cellular and tissue tropism can influence pathogenicity by expanding the range of susceptible cell types and target organs, and by promoting infection of cells within the lower respiratory tract, all of which can increase disease severity. Cellular tropism is in part dictated by receptor and co-factor usage of which HCoVs use a variety of receptors including aminopeptidase N (APN), dipeptidyl peptidase 4 (DPP4), transmembrane serine protease 2 (TMPRSS2), angiotensin converting enzyme 2 (ACE2) and 9-O-acetylated sialic acids (*3, 15–18*). Although HCoV-NL63, SARS-CoV and SARS-CoV-2 all utilise the same receptor (ACE2), they have vastly different CFRs and clinical disease outcomes indicating that receptor usage alone cannot explain pathogenicity.

Several studies have shown that SARS-CoV, MERS-CoV and SARS-CoV-2 can all replicate efficiently in the lower respiratory tract affecting alveoli of humans, ex vivo lung tissue and human alveolar type II cultures (*19, 20*). In contrast, replication of seasonal HCoVs appears to be limited to the upper respiratory tract in the majority of cases and show lower replication in human alveolar type II cultures (*21, 22*). However, the exact requirements for HCoV infection within different anatomical sites of the respiratory tract is unknown. Systematic analysis of the host cellular response, particularly with SARS-CoV-2, has uncovered a plethora of cellular pathways induced in response to infection, antagonised by viral accessory proteins or harnessed by virus to facilitate productive viral replication (*23–26*). Several, including the MAPK and NF-κB pathways, have been shown to be broadly associated with other HCoV strains (*23*). Critically, many of these analyses were conducted in immortalised cell lines but comparative studies of pathogenic and seasonal HCoVs in human airway tissue are important as virus-host interactions that determine tropism, replication kinetics, innate immune activation and tissue damage are highly context dependent.

Here, we used physiologically relevant in vitro models of the human respiratory tract to dissect virus-intrinsic and host-mediated determinants of HCoV pathogenicity. We found that pathogenic HCoVs replicate to higher levels in lung alveolar type II (AT2) cells and induce higher levels of type III interferons compared to other anatomical sites in the airways. HCoV-229E and MERS-CoV both caused significant cell death compared to the other HCoVs in AT2 cells. Gene expression analysis in the nasal epithelium and AT2 cells identified several host pathways associated with infection by pathogenic viruses as well as divergent responses to HCoV-229E when compared to other seasonal HCoVs. These data expand on the complexity of interactions between the host and HCoVs in the respiratory tract.

## Results

### Pathogenic HCoVs replicate to higher titres in lung AT2 cells compared to seasonal HCoVs

To compare respiratory tract tropism and replication capacity of different pathogenic and seasonal HCoVs, we inoculated primary human nasal epithelial cells (HNEC), immortalised bronchial cells (BCi) and small airway epithelial cells (SABCi) differentiated at an air-liquid interface (ALI) along with embryonic stem cell-derived alveolar type II (edAT2) cells representing cells of the human airway from the nose, trachea, small airway and lung alveoli (*27*). Viruses compared include two SARS-CoV-2 isolates (ancestral (B.1) and BA.5 variants), MERS-CoV, HCoV-229E, HCoV-NL63 and HCoV-OC43, where infectious virus titre (or viral RNA for HCoV-OC43) was quantified in the supernatant (apical and basal for ALI) over 7 days (Fig. 1). No virus was detected in the basal medium of virus-infected ALI cultures (data not shown).

**Figure 1:**
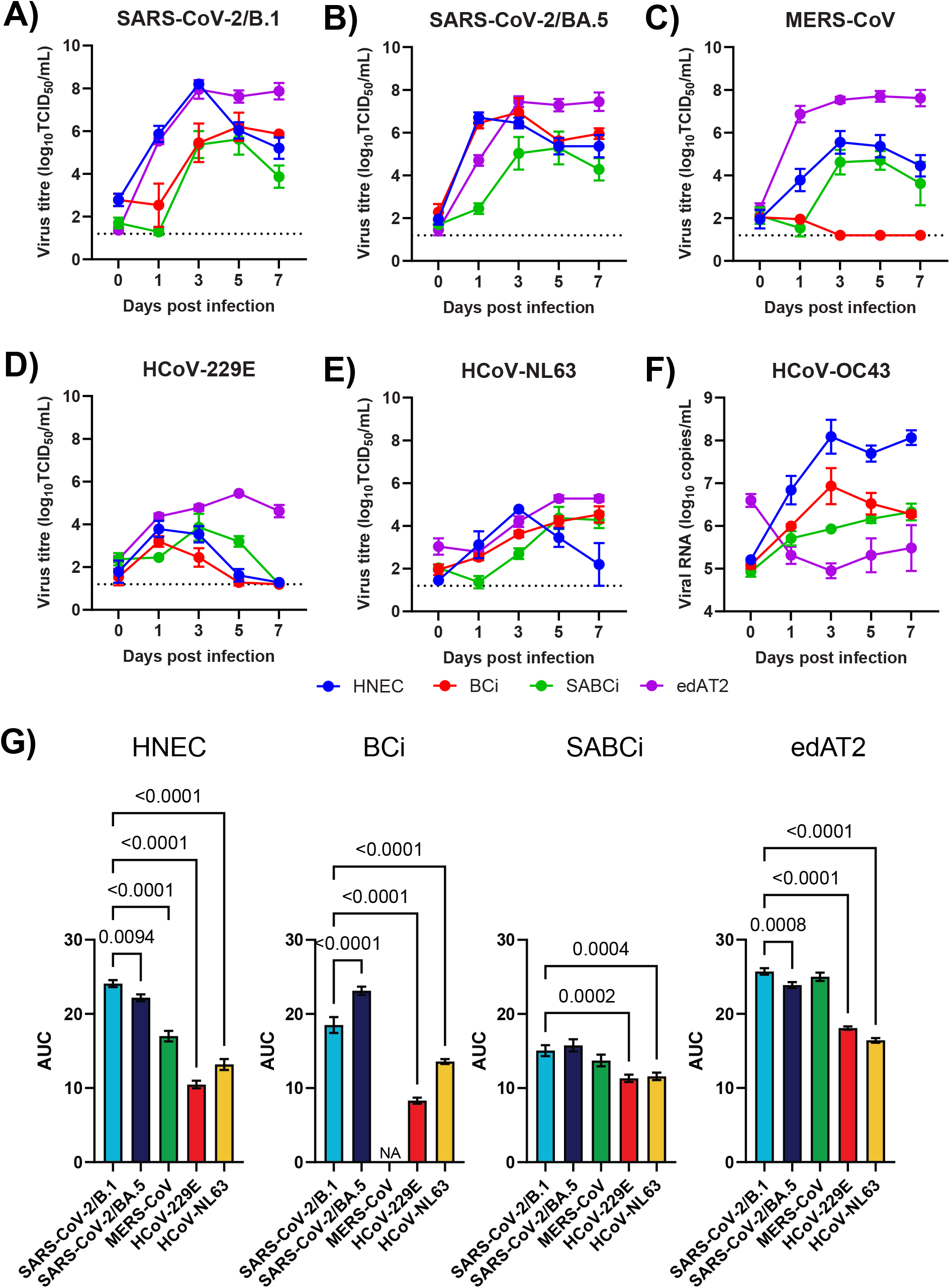
Pathogenic HCoVs replicate to higher titres in lung epithelial cells compared to seasonal HCoVs. Infectious virus titres in HNECs (blue), BCi (red), SABCi (green) and edAT2 (purple) cultures infected with SARS-CoV-2/B.1 I **(A)**, SARS-CoV-2/BA.5 **(B)**, MERS-CoV **(C)**, HCoV-229E **(D)** and HCoV-NL63 **(E)** shown as log10 50% tissue culture infectious units per ml (TCID50/ml). **(F)** Viral genome copies in HNECs (blue), BCi (red), SABCi (green) and edAT2 (purple) cultures infected with HCoV-OC43 represented as log10 genome copies per ml. Data are representative of two experimental repeats, each with three replicates. Mean ± SD is shown. Dotted lines represent lower limit of detection. **(G)** Area under the curve analysis in HNECs, BCi, SABCi and edAT2 cells using data from **(A)** to **(E)**.

SARS-CoV-2 variants replicated to high titres across all four cell model systems (Fig. 1A, B). BA.5 reached its peak titre faster than B.1, although the peak was lower than B.1 (6.7 log10 TCID50/ml at 1-day post-infection (dpi) compared to 8.2 log10 TCID50/ml at 3 dpi) in HNECs. While BA.5 demonstrated faster replication in BCi cultures compared to B.1 (6.5 versus 2.5 log10 TCID50/ml at 1 dpi), there was no significant difference in peak titres (7.0 at 3 dpi for BA.5 compared to 6.2 log10 TCID50/ml for B.1 at 5 dpi). B.1 and BA.5 showed similar replication kinetics in SABCi and edAT2 cells, reaching peak titres of 5.6 and 5.3 log10 TCID50/ml in SABCi, respectively and 7.8 and 7.7 log10 TCID50/ml in edAT2, respectively. MERS-CoV showed robust replication in HNECs, with a peak viral titre of 6.4 log10 TCID50/ml on 3 dpi (Fig. 1C). Notably, MERS-CoV did not replicate in BCi cultures but did replicate in primary bronchial epithelial cell cultures differentiated from three donors (Sup. Fig. 1A). MERS-CoV showed similar replication kinetics in SABCi and edAT2 cells as SARS-CoV-2, with peak titres of 4.7 and 7.7 log10 TCID50/ml, respectively at 5 dpi.

HCoV-229E showed modest replication in HNEC, BCi and SABCi, reaching peak titres of 4.0, 3.2 and 3.7 log10 TCID50/ml, respectively. Unlike the other viruses, infectious HCoV-229E was undetectable at 7 dpi (Fig. 1D). Lower virus titers in airway cultures appears to be unique to HCoV-229E as has been observed previously (*21*). HCoV-NL63 also showed modest replication across HNEC, BCi and SABCi cultures, reaching peak titres of 4.8, 4.5 and 4.4 log10 TCID50/ml, respectively (Fig. 1E). In edAT2 cells, both HCoV-229E and HCoV-NL63 replicated to their highest levels peaking at 5 dpi at 5.5 and 5.3 log10 TCID50/ml (Fig. 1D, E). To study OC43 replication in the respiratory tract, we selected a clinical isolate as we have previously shown that the reference VR-1558 virus does not infect human airway cultures (*21*). We used qPCR of apical wash samples as a readout for virus production because we have not identified an immortalised cell line in which HCoV-OC43 clinical isolates replicate. We found HCoV-OC43 replicated well in HNECs, reaching 8.3 log10 genome copies/ml at 3 dpi, but only modest replication in BCi (peak of 6.9 log10 copies/ml at 3 dpi) and SABCi (peak of 6.3 log10 copies/ml at 7 dpi) (Fig. 1F). Despite high levels of OC43 in edAT2 cells on 0 dpi (6.6 log10 copies/ml), no replication was observed (Fig. 1F). This is in contrast with the reference strain for OC43 VR-1558 which replicates robustly in edAT2 cells (*21*).

SARS-CoV-2/B.1 and HCoV-OC43 reached their highest titres in HNECs representative of the upper respiratory tract, while the remaining viruses reached their highest titres in edAT2 cells representative of the lower respiratory tract (Fig. 1). Infectious titres were higher for SARS-CoV-2 and MERS-CoV compared to HCoV-NL63 and HCoV-229E across all airway models (Fig. 1G). In HNECs, BCi and SABCi MERS-CoVs replication was lower than SARS-CoV-2. However, in edAT2 cells there was clear segregation of virus titre levels between pathogenic (SARS-CoV-2 and MERS-CoV) and seasonal (HCoV-229E and HCoV-NL63) HCoVs (Fig. 1G). This suggests that the ability of particular HCoVs to replicate to high titres in edAT2 cells may distinguish pathogenic from milder seasonal HCoVs.

### Pathogenic HCoVs induce higher type III interferon secretion compared to seasonal HCoVs across the airways

To investigate the host epithelial response following infection, we measured the secretion of 13 cytokines at 0, 2, 4 and 6 dpi in each of the airway cultures using a cytokine bead array. Overall, these data indicated that there was a gradient of cytokine expression across the respiratory tract with the highest cytokine expression following HCoV infection observed in HNECs and the lowest in edAT2 cells (Fig. 2A). Of the 13 cytokines, these airway models predominantly expressed IP-10 and interferons (IFNs) in response to HCoV infection. At 2 dpi, a 2.5-fold to 30-fold increase in expression of CXCL10 was observed following infection with HCoV-OC43 and SARS-CoV-2 in HNECs. All viruses increased CXCL10 expression by HNECs up to approximately 90-fold over the first 4 dpi. CXCL10 was also increased 80-fold on 4 dpi by BA.5 infection and 6 to 8-fold by HCoV-229E and HCoV-NL63 on 6 dpi in BCi as well as up to 26-fold by HCoV-229E, B.1 and BA.5 in SABCi and up to 2.5-fold by HCoV-229E and HCoV-NL63 in edAT2 cells. Overall, all HCoVs strongly induced IP-10 secretion in HNECs, with HCoV-229E showing the broadest expression across the airways.

**Figure 2:**
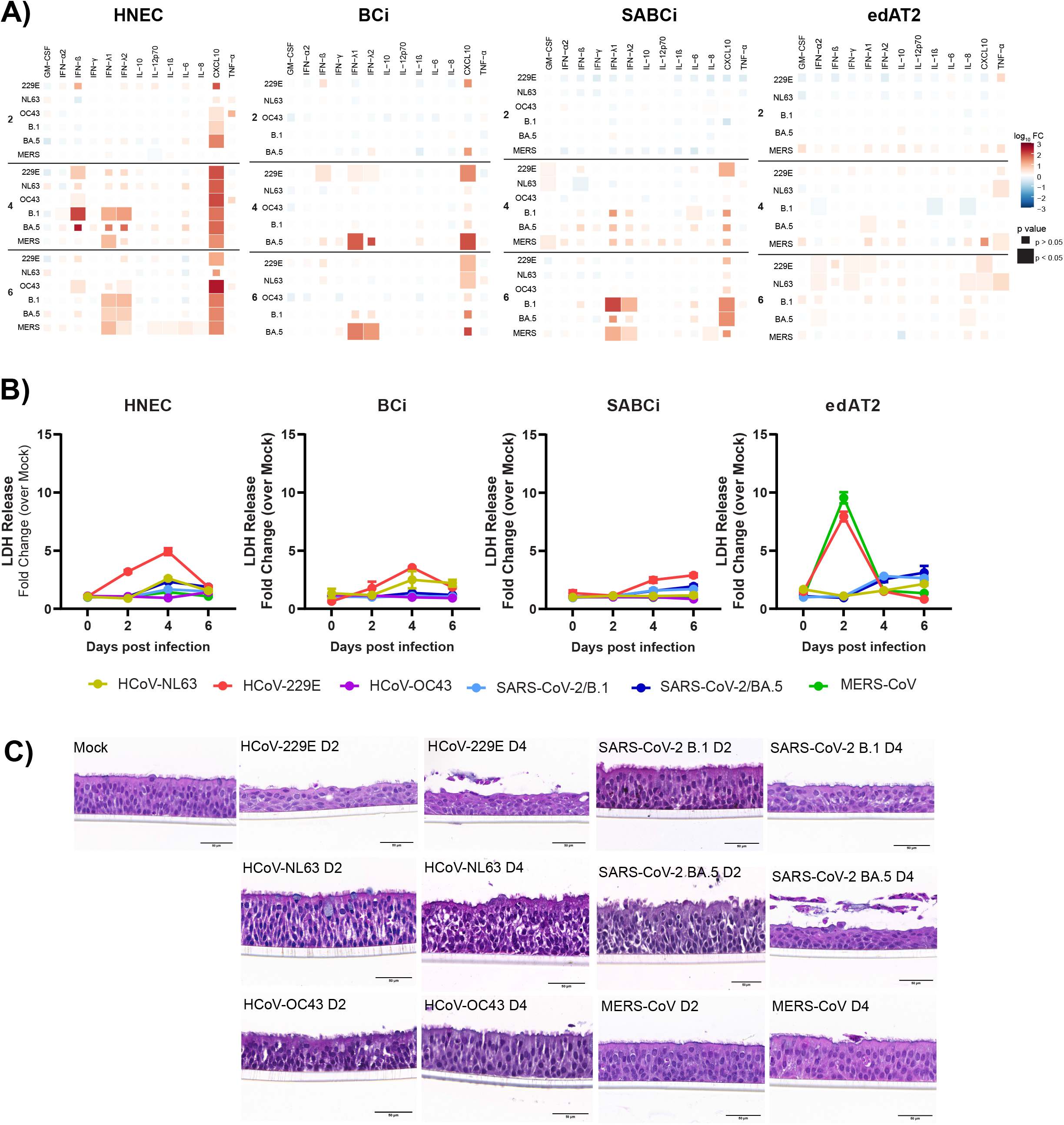
HCoV-229E causes increased epithelial cell damage and death compared to other HCoVs across the airways. **(A)** Heatmap showing cytokine expression represented as log10 fold change relative to mock controls across HNEC, BCi, SABCi and edAT2 cells infected with HCoV-229E, HCoV-NL63, HCoV-OC43, SARS-CoV-2/B.1, SARS-CoV-2/BA.5 and MERS-CoV. Statistical comparisons were performed using a student t-test. **(B)** Lactate dehydrogenase (LDH) levels in the basal medium of HNEC, BCi and SABCi cultures or in the supernatant of edAT2 cells following infection with HCoV-NL63 (yellow), HCoV-229E, (red), HCoV-OC43 (purple), SARS-CoV-2/B.1 (light blue), SARS-CoV-2/BA.5 (dark blue) and MERS-CoV (green). LDH release is shown as a fold change compared to mock controls. Data are representative of two experimental repeats, each with three replicates. Mean ± SD is shown. **(C)** Representative images of HNEC sections infected with HCoV-229E, HCoV-NL63, HCoV-OC43, SARS-CoV-2/B.1, SARS-CoV-2/BA.5 and MERS-CoV fixed at 2 or 4 dpi and stained with haematoxylin and eosin. Scale bar = 50 µm.

SARS-CoV-2 and MERS-CoV infection significantly increased IFN-λ1 and IFN-λ2 approximately 1.2 to 8.7-fold and 2.3 to 18-fold in HNECs, respectively. HCoV-OC43 infection also increased IFN-λ2 secretion on 6 dpi by 1.7-fold. In contrast, HCoV-229E and HCoV-OC43 modestly increased IFN-β approximately 1.6 to 12.5-fold compared to SARS-CoV-2/B.1 that increased IFN-β 100-fold on 4 dpi. SARS-CoV-2/B.1 also increased IFN-α by 1.3-fold on 4 dpi. In BCi, SARS-CoV/BA.5 induced a >19-fold increase in IFN-λ1 and IFN-λ2 at 6 dpi while only modest expression of IFN-β (2.6-fold) and IFN-λ1/2 (1.5-1.7-fold) was induced by HCoV-229E at 4 dpi. In SABCi, only B.1 and MERS-CoV increased IFN-λ1/2 (3.8 to 100-fold) at 6 dpi. In edAT2 cells, low expression of IFN-λ1 and IFN-α was induced following BA.5 infection (<1.7-fold) as well as IFN-α, IFN-λ1 and IFN-γ following HCoV-229E (<3.2-fold) and HCoV-NL63 (<1.6-fold) infection. Overall, these data indicate that IFN-λ was expressed at higher levels following infection with pathogenic HCoVs compared to seasonal HCoVs.

Higher virus titres may partially explain this difference in the upper airways but this did not extend to edAT2 cells.

### HCoV-229E causes cytotoxicity in HNECs and edAT2 cells despite modest replication

To assess cell viability following infection, we measured the levels of lactate dehydrogenase (LDH) in the supernatant (basal for ALI cultures) on 0, 2, 4 and 6 dpi. As shown in Figure 2B, cell death was minimal in HNEC, BCi and SABCi cultures with only HCoV-229E increasing the levels of LDH relative to mock-inoculated cells in HNEC at 4 dpi (5-fold), BCi at 4 dpi (3.5-fold) and SABCi at 6 dpi (2.5-fold). However, both HCoV-229E and MERS-CoV led to an approximately 8-10-fold increase in the levels of released LDH, indicative of significant cell death in edAT2 cells at 2 dpi (Fig. 2B). This was associated with detection of observable CPE by light microscopy (data not shown). This suggests that MERS-CoV and HCoV-229E are causing changes in cell viability in edAT2 cells while HCoV-229E caused more cell death than the other HCoVs across the airways.

To further explore tissue integrity of the ALI cultures, we stained cross-sections of mock and HCoV-infected HNEC, BCi and SABCi ALI cultures with haematoxylin and eosin (H&E) at 2 and 4 dpi (Fig. 2C, Sup Fig. 1B). No obvious changes in cell morphology or ciliation levels were observed in HNECs infected with HCoV-OC43, HCoV-NL63 or MERS-CoV at either time point (Fig. 2C). Following SARS-CoV-2/B.1 infection there was a modest reduction in ciliated cells and flattening of the stratified epithelium at 4 dpi. In contrast, HNECs infected with SARS-CoV-2/BA.5 at 4 dpi and HCoV-229E at 2 and 4 dpi showed extensive loss of apical cilia, a flattened, squamoid cellular morphology and evidence of cell shedding which are suggestive of reduced tissue integrity and potential cellular dedifferentiation. No obvious changes in cellular morphology or epithelium integrity were observed following HCoV infection in BCi or SABCi cells (Sup Fig. 1B, C). These data are consistent with LDH release and indicate that HCoV-229E causes the most epithelial damage of all the HCoV strains in HNECs. Given that the differences between pathogenic and seasonal HCoVs were predominately observed in HNECs and edAT2 cells we focused on these two cell models for the rest of the study.

### HCoV-229E and SARS-CoV-2 induce dramatic shifts in responding nasal epithelial cell phenotypes

To identify host-specific transcriptional changes that occur following HCoV infection in the nasal epithelium, we performed single cell RNA sequencing (scRNA-seq) on HNECs at 1- and 3-dpi following SARS-CoV-2 (B.1 and BA.5), MERS-CoV, HCoV- 229E, HCoV-NL63 and HCoV-OC43 infection (Fig. 3A). Following sample filtering and integration of the data, we analysed the transcriptomes of 21,689 cells. We identified 16 epithelial clusters and designated epithelial cell types based on gene expression clustering and known markers of epithelial differentiation (Fig. 3B, C, D). As shown in Figure 3E and Sup. Fig 2, these included basal cells (TP63hi and KRT5hi), secretory cells (MSMB+ and CYP2F1+), ciliated cells (FOXJ1+, CDHR3+ and DNAH9+) and ionocytes (FOXI1+ and CFTRhi). Basal cells were further separated into cycling basal cells (MKI67hi), basal 1, basal 2 and suprabasal. Secretory cells were separated into club, goblet 1 and goblet 2 populations. In addition, we identified a cluster of cells that expressed high levels of basal (KRT5hi) and club (CYP2F1+) markers that were denoted ‘basal-club’ cells and deuterosomal (DEUP1+and FOXN4+) cell populations. In samples that were exposed to viral infection, we identified an additional four cell types; a ciliated cell cluster that showed high expression of the unfolded protein response gene DNA damage-inducible transcript 3 (DDIT3, Sup Fig. 2), denoted DDIT3hi ciliated cells, followed by four clusters of cells that showed high expression of Interferon-stimulated gene 15 (ISG15) and other antiviral proteins that were denoted responding basal, responding basal-club, responding ciliated and responding secretory cells based on expression of the ciliated, secretory and basal markers above (Fig. 3E).

**Figure 3:**
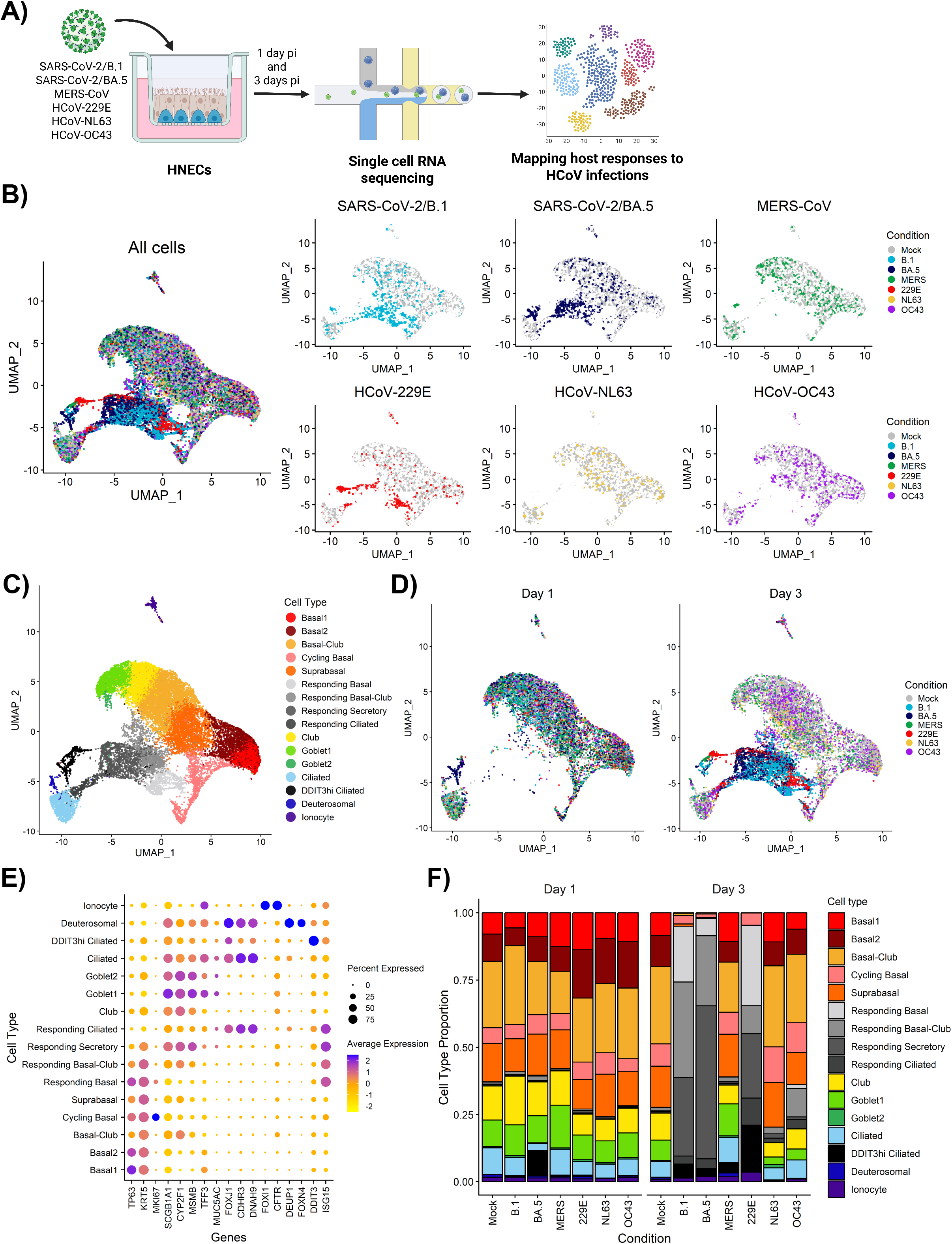
Single cell RNA sequencing of HCoV-infected HNECs shows that SARS-CoV-2 and HCoV-229E drastically increase responding cell populations. **(A)** Schematic representation of the experiment performed. HNECs were inoculated with SARS-CoV-2/B.1, SARS-CoV-2/BA.5, MERS-CoV, HCoV-229E, HCoV-NL63 or HCoV-OC43. After 1 or 3 dpi, cells were dissociated and prepared for single cell RNA sequencing. **(B)** Uniform Manifold Approximation and Projection (UMAP) representation of cell profiles from each experimental condition. Dots represent individual cells from each condition; grey: mock, light blue: SARS-CoV-2/B.1 (B.1), dark blue: SARS-CoV-2/BA.5 (BA.5), green: MERS-CoV (MERS), red: HCoV-229E (229E), yellow: HCoV-NL63 (NL63), purple: HCoV-OC43 (OC43). **(C)** UMAP representation of the cell profiles from all experimental conditions with epithelial cell types annotated. **(D)** UMAP representation of cell profiles from each experimental condition stratified by time point (1 and 3 dpi). **(E)** Dot plot of marker genes for distinct cell types. Colour scale indicates the mean of normalized expression of marker genes in each cell type and dot size is proportional to the percentage of cells within each cell cluster expressing the marker genes. **(F)** Proportion of each defined cell type across experimental conditions and time-points.

Next, we compared the relative proportion of each cell type across virus infection conditions and time-points (Fig. 3F). Mock-infected samples showed 24-28.3% basal cell types (including basal 1, basal 2 and cycling basal), 14.2-15.3% suprabasal, 24.6-28.7% basal-club,10.2-12.6% club, 7.9-10.5% goblet cell types (goblet 1 and goblet 2), 6.4-10.4% ciliated cells, 0.6-1% deuterosomal and 0.8-1.6% ionocytes in 1- and 3 dpi samples. The proportion of cell types were similar to mock-infected samples at 1 dpi for SARS-CoV-2/B.1, MERS-CoV, HCoV-229E, HCoV-OC43 and HCoV-NL63.

However, SARS-CoV-2/BA.5 showed a population of DDIT3hi ciliated cells (9.4%) which was absent in mock-infected cells. At 3 dpi, the proportion of DDIT3hi ciliated cells increased to 4.6%, 2.5%, 3.9% and 17.3% for SARS-CoV-2/B.1, SARS-CoV-2/BA.5, MERS-CoV and HCoV-22E, respectively. HCoV-OC43 and HCoV-NL63 showed similar cell type proportions to mock-infected cells, including no change in the proportion of DDIT3hi ciliated cells. In addition, we observed drastic shifts in the proportion of epithelial cell types for HCoV-229E and SARS-CoV-2 (B.1 and BA.5), dominated by the responding basal, responding basal-club, responding secretory and responding ciliated populations. MERS-CoV, HCoV-NL63 or HCoV-OC43 did not induce a drastic shift in responding cell populations.

Analysis of cell cycle indicated that the frequency of cells in G1 phase following infection with SARS-CoV-2 (B.1 and BA.5) and HCoV-229E on 3 dpi was increased compared with the mock condition (Sup. Fig. 3A-C). This change was likely associated with the appearance of responding cells which were predominantly in G1 phase. To determine the source of the responding and DDIT3hi cell types we preformed RNA velocity analysis (*28*). While there were small differences depending on the virus, ciliated cells were the most likely source for the DDIT3hi population for HCoV-229E, SARS-CoV-2 (B.1 and BA.5) and MERS-CoV (Sup. Fig. 4). For HCoV-229E, SARS-CoV-2/B.1 and SARS-CoV-2/BA.5 the data suggested that basal cycling cells were the source of the responding basal population (Sup. Fig. 4). Basal, suprabasal and basal-club cells all lead to the responding basal-club population. Secretory and club cells were the source for responding secretory cells and ciliated and DDIT3hi ciliated cells were the source for responding ciliated cells.

**Figure 4:**
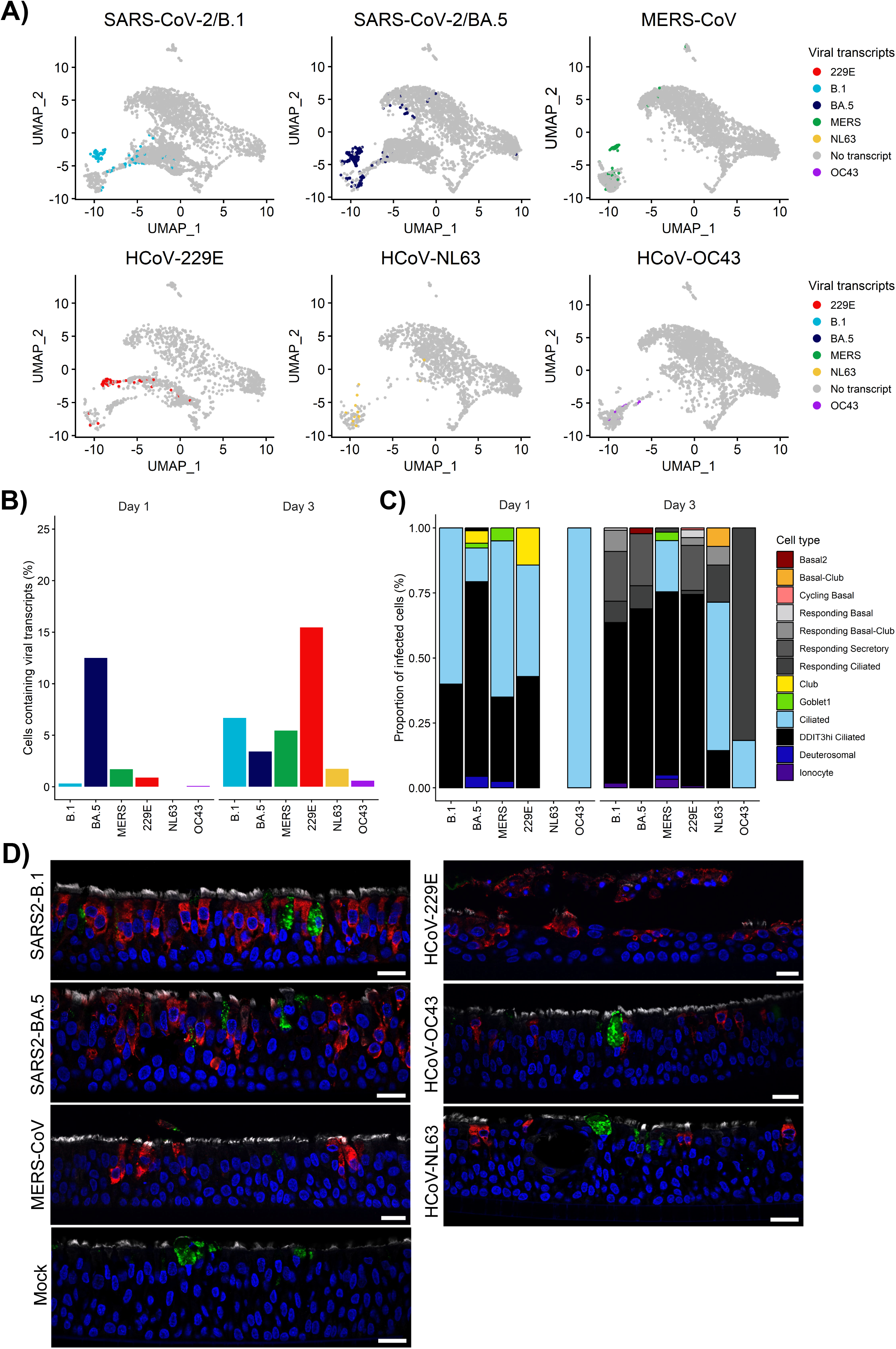
HCoVs show preferential tropism for ciliated epithelial cells. **A)** UMAP representation of cell profiles from each experimental condition (other than mock). Grey dots represent cells containing no viral transcripts and coloured dots represent cells with detectable viral transcripts: light blue: SARS-CoV-2/B.1 (B.1), dark blue: SARS-CoV-2/BA.5 (BA.5), green: MERS-CoV (MERS), red: HCoV-229E (229E), yellow: HCoV-NL63 (NL63), purple: HCoV-OC43 (OC43). **(B)** Bar plots of the relative proportion of infected cells across time points. The x axis represents the different viruses, and the y axis indicates proportion of cells containing transcripts. **(C)** Bar plots of the relative proportion of cell types containing viral transcripts across time points. The x axis represents the different viruses, and the y axis indicates the proportion of each cell type, with the total proportions of all cell types summing to 100% per virus on 1 and 3 dpi. **(D)** Representative immunohistochemistry stain of HNECs fixed at 2 dpi following virus infection. Sections were immunostained to identify infected cells (viral nucleocapsid, red), ciliated cells (α-Tubulin, white), and or goblet cells (Mucin 5AC [MUC5AC], green). Sections were counter stained with DAPI (blue). Images are representative of two independent experiments. Scale bar = 20 μm.

### Pathogenic and seasonal HCoV predominantly infect ciliated epithelial cells but only cells infected with HCoV-229E lose their ciliated cell identity

To determine the cellular targets of HCoVs in the airways, we first determined the proportion of all cells containing viral transcripts in HNECs following exposure to each virus. On 1 dpi, infected cells were observed for SARS-CoV-2 (B.1 and BA.5), MERS-CoV, HCoV-229E and HCoV-OC43 but the highest percentage of infected cells were those infected with BA.5 (Fig. 4A). Most of these cells were either ciliated or ciliated cells expressing increased DDIT3 indicative of cellular damage (Fig. 4B). On day 3, viral transcripts were detected for all viruses with the highest proportions observed for HCoV-229E (15.5%), SARS-CoV-2/B.1 (6.7%) and MERS-CoV (5.5%, Fig. 4A-B).

Most of these cells were either ciliated, responding ciliated or DDIT3hi ciliated cells (Fig. 4C). A fraction of SARS-CoV-2 (B.1 and BA.5) and HCoV-229E-infected cells were responding secretory cells.

To confirm this finding, we stained for viral nucleocapsid protein along with markers for goblet cells (MUC5AC) and ciliated cells (α-Tubulin) in HNECs fixed at 2 dpi. Infected cells were observed adjacent to α-Tubulin for all HCoVs (Fig. 4D) confirming that virus was located primarily within the cytoplasm of ciliated cells. For SARS-CoV-2/BA.5 and HCoV-229E, some nucleocapsid protein was associated with non-α Tubulin, non-MUC5AC-expressing cells. Overall, these data show that ciliated cells are the major target of HCoVs in HNECs but that a low percentage of other, non-ciliated cell types can be infected. Similar to our H&E analysis, the expression of α-Tubulin was reduced following infection with HCoV-229E in HNECs indicative of loss of cilia (Fig. 4D). To determine if this tropism was also observed in the large and small airways, we stained for virus and cell markers in BCi and SABCi at 4 dpi. Overall, fewer nucleocapsid positive cells were observed for all HCoVs in BCi and SABCi cultures (Supp. Fig. 5A-B). However, all infected cells co-expressed α-Tubulin, indicative of infection of ciliated cells. This confirms that ciliated cells are the major cell target for HCoVs across the airways.

**Figure 5:**
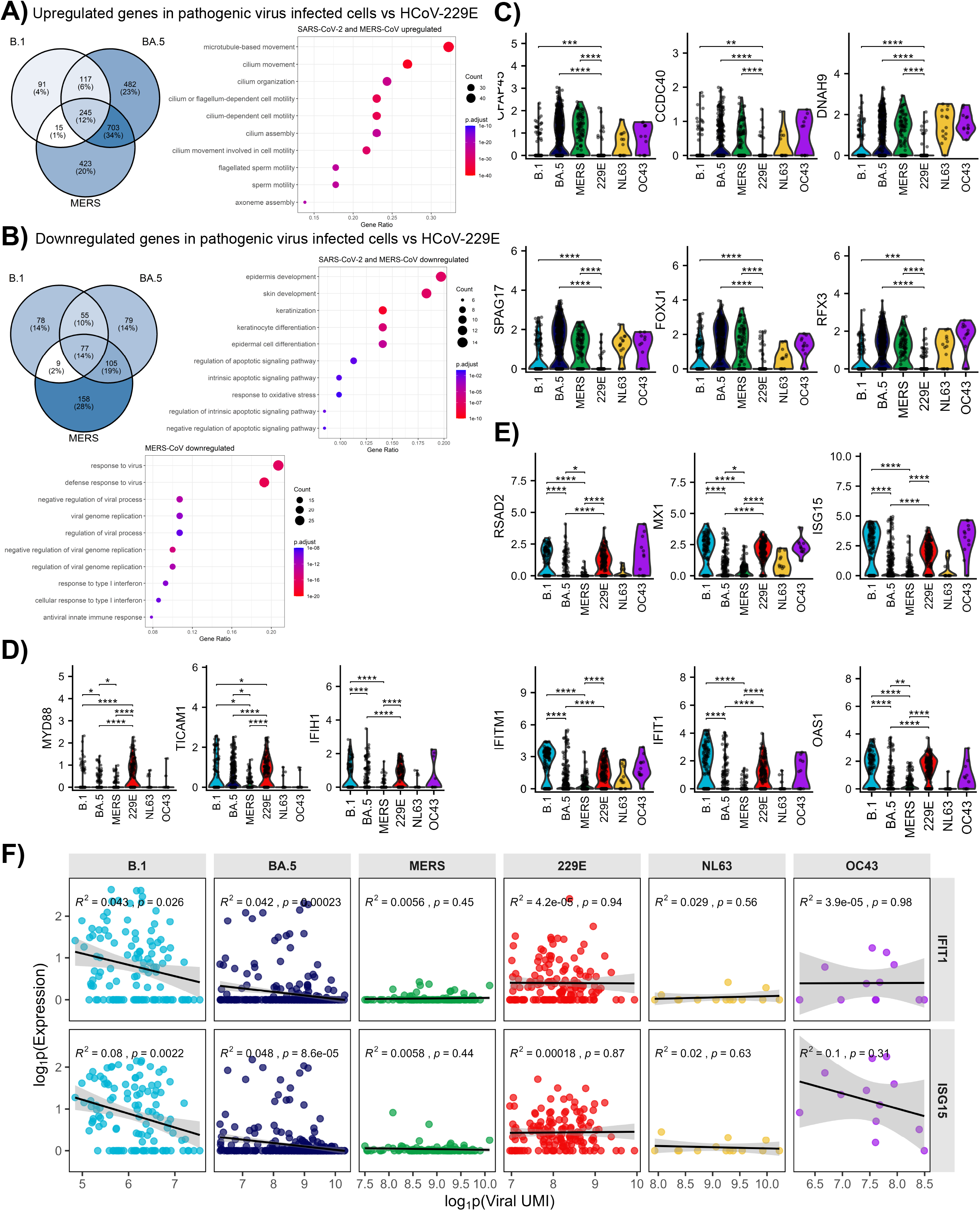
HNECs infected with HCoV-229E shows signs of reduced ciliogenesis compared to other HCoVs. **(A)** Venn diagram of the number (and percentage) of genes with up-regulated expression and top 10 Gene Ontology Biological Pathways enriched in transcript-containing cells following infection with pathogenic HCoVs (B.1, BA.5, MERS-CoV) compared to HCoV-229E. **(B)** Venn diagram of the number (and percentage) of genes with down-regulated expression and top 10 Gene Ontology Biological Pathways enriched in transcript-containing cells following infection with pathogenic HCoVs (B.1, BA.5, MERS-CoV) compared to HCoV-229E. Colour indicates adjusted p-value and size of dot indicates number of genes in each pathway. Expression level of genes associated with ciliated cell function **(C)**, viral detection **(D)** and interferon signalling **(E)** in cells containing viral transcripts across both 1 and 3 dpi. **(F)** Linear regression analysis of viral gene expression (log1p(Viral UMI) and IFIT1 (top) and ISG15 (bottom) expression (log1p(expression) for each cell containing viral transcripts split by virus infection. Black lines show the slope and grey area shows the 95% confidence interval. Values represent the R2 and p value calculated by Pearson correlation.

We next evaluated the expression of entry receptors in HNECs across all conditions (Supp. Fig. 6A). Overall, only low proportions of all cell-types expressed ACE2 and DDP4 with expression primarily detected in responding and secretory cell populations. APN and TMPRSS2 were expressed primarily in ciliated and secretory cells. CTSL was expressed across all cell types (Supp. Fig. 6B). Interestingly, DDIT3hi expressing cells and cells containing viral transcripts had low expression of all known HCoV entry receptors suggesting that infection may downregulate the expression of these receptors.

**Figure 6:**
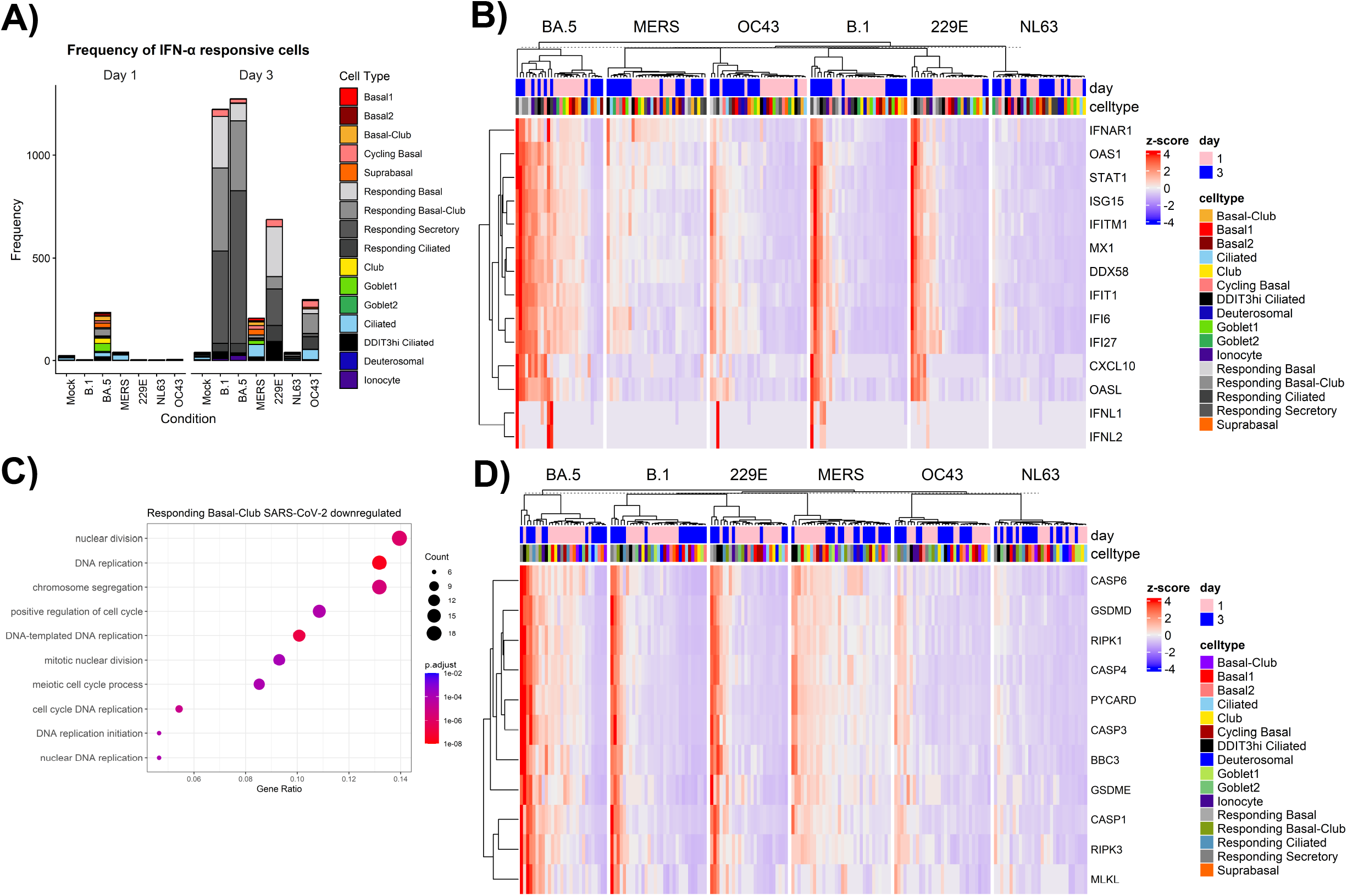
Responding Cells are distinguished by high expression of interferon-stimulated genes. **(A)** Frequency of IFNα responsive cells from each experimental condition on 1 and 3 dpi. **(B)** Z-score of selected genes associated with interferon signalling in each condition, cell type and day post infection. **(C)** Top 10 Gene Ontology Biological Pathways enriched in responding basal-club cells following HCoV-229E infection compared with SARS-CoV-2 (B.1, BA.5). Colour indicates adjusted p-value and size of dot indicates number of genes in each pathway. **(D)** Z-score of selected genes associated with cell death in each condition, cell type and day post infection.

Given the low frequency of cells infected with either HCoV-OC43 or HCoV-NL63, we focused our comprehensive evaluation of infected cells on SARS-CoV-2 (B.1 and BA.5), HCoV-229E and MERS-CoV at 1 and 3 dpi. Overall, 245 genes (11.8% of total genes up-regulated by any pathogenic HCoV) were up-regulated and 77 genes (13.5% of total genes up-regulated by any pathogenic HCoV) were down-regulated by all pathogenic viruses (SARS-CoV-2 and MERS-CoV) compared to HCoV-229E (Fig. 5A, B). Pathway analysis indicated that cells infected with the pathogenic HCoVs had higher expression of genes associated with ciliated cell function, including cilia assembly and movement compared to HCoV-229E (Fig. 5B). Conversely, infection with HCoV-229E induced higher expression of genes associated with epithelial differentiation compared to pathogenic HCoVs. To further explore how virus infection changed ciliated cell identity, we compared all cells containing virus transcript (of which the majority are ciliated cells) to all ciliated cells from the mock condition. Overall, HCoV-229E reduced the expression of several ciliated cell identity genes (DNAH9, CCDC40, CFAP45, FOXJ1 and SPAG17) compared to other viruses and uninfected ciliated cells (Fig. 5C). Taken together these data suggest that HCoV-229E affects epithelial cell differentiation leading to a reduction in ciliated cell function more so than pathogenic HCoVs. Interestingly, cells containing HCoV-229E transcripts were also predominantly in G1 phase, which contrasted with all other infected cells which were in S phase (Sup. Fig. 3C). This suggests that infection with HCoV-229E in ciliated cells leads to a distinct cellular response compared to other HCoVs.

Pathway analysis of pathogenic viruses compared to HCoV-229E also showed that only MERS-CoV infected cells demonstrated reduced expression of genes associated with innate immune responses to viral infection (Fig. 5B). Further investigation of the expression of genes associated with the innate immune response revealed that MERS-CoV and SARS-CoV-2/BA.5 demonstrated reduced expression of IFIH1 and TICAM1, key genes associated with MDA5- and TRIF-dependent RNA-sensing pathways (Fig. 5D). In addition, MERS-CoV and SARS-CoV-2/BA.5 infected cells demonstrated reduced expression of interferon stimulated genes (ISGs), including IFIT1, IFITM1, ISG15, MX1, RSAD2 and OAS1 compared to SARS-CoV-2/B.1 and HCoV-229E infected cells (Fig. 5E). For SARS-CoV-2 (B.1 and BA.5) we found IFIT1 and ISG15 expression in infected cells negatively correlated with the number of viral transcripts per cell (Fig. 5F). For MERS-CoV, the levels of IFIT1 and ISG15 were consistently low regardless of viral transcript level. In contrast, we did not find this for HCoV-229E, HCoV-OC43 or HCoV-NL63, although the latter two were limited by low infected cell numbers. Overall, these data suggest that pathogenic HCoVs can dampen the antiviral innate immune response in infected HNECs, with SARS-CoV-2/BA.5 and MERS-CoV being more efficient at this than SARS-CoV-2/B.1.

### Cells responding to HCoV infections are defined by a strong IFN signature

Next, we evaluated the bystander cells responding to virus infection. These cell populations, which were observed primarily following SARS-CoV-2, HCoV-229E and to a lesser extent HCoV-OC43 infection, were distinguished by a high expression of ISGs (Fig. 6A and Sup. Fig 7A). Interestingly, MERS-CoV infection also increased the number of cells with IFN signalling, but these were not ‘responding’ populations but rather a mix of different cell types like that observed for BA.5 at 1 dpi (Fig. 6A). Limited ISG expression was observed following HCoV-NL63 infection, while SARS-CoV-2 BA.5 infection led broad ISG induction across cell types (Fig. 6B and Sup. Fig 7B). Comparison of responding cells following SARS-CoV-2 and HCoV-229E infection suggested that irrespective of virus these cells show signatures predominantly associated with responses to stimulus in addition to metabolic changes (Sup Fig. 8). Interesting, responding basal-club cells following HCoV-229E infection showed an induction of genes associated with DNA replication and cellular division suggesting that these cells may have a different phenotype compared to responding basal-club cells induced following SARS-CoV-2 infection (Fig. 6C).

**Figure 7:**
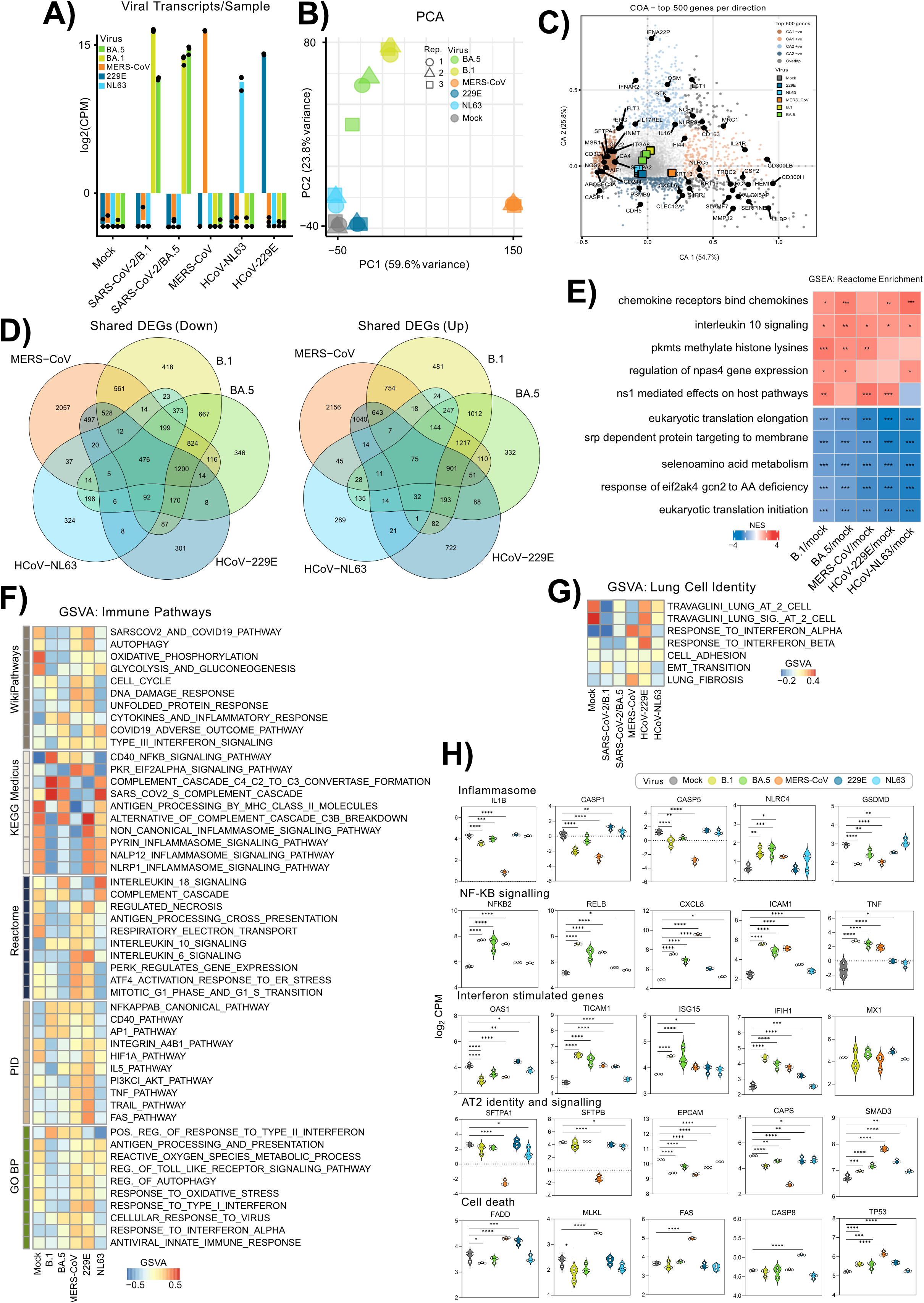
HCoVs induced fibrotic signatures in edAT2 cells but only HCoV-229E and MERS-CoV cause cell death. **(A)** Viral gene expression in edAT2 cells (log2 counts per million (CPM)) at 3 dpi (3 replicates each), in each cell condition (Mock, B.1, BA.5, MERS-CoV, HCoV-229E and HCoV-NL63). **(B)** PCA of all samples in each condition. **(C)** Correspondence analysis (COA) of all AT2 cell conditions with top 500 genes highlighted in each direction (positive or negative) along the top 2 COA axes (CA1, CA2). **(D)** Venn diagram of the number of genes with up-regulated or down-regulated expression in each condition against mock. **(E)** Gene set enrichment analysis showing the top shared Reactome based gene sets across each of the 5 DGE analyses (each viral infection against mock) *P adj. < 0.05, ** P adj. < 0.01, P adj. < 0.001. Gene set variation analysis (GSVA) scores associated with lung cell identity **(F)** and top immune pathways **(G)** across all conditions. **(H)** Gene expression signatures of inflammasome, NF-kB signalling, interferon stimulated genes, AT2 identity and signalling and cell death associated genes across all conditions. FDR is shown. * < 0.05, ** < 0.01, *** < 0.001, **** < 0.0001

### SARS-CoV-2 and HCoV-229E infection of HNECS induces strong cell death signatures in infected and bystander cells

Given that we observed some cell death in HNECs following HCoV-229E and SARS-CoV-2/BA.5 infection (Fig. 2), we examined gene expression associated with cell death pathways. MERS-CoV increased cell death gene expression predominantly in DDIT3hi ciliated cells on both 1 and 3 dpi (Fig. 6D). SARS-CoV-2/B.1, SARS-CoV-2/BA.5 and HCoV-229E induced cell death in DDIT3hi cells and responding cells on 3 dpi with SARS-CoV-2/BA.5 also increasing these signatures on 1 dpi. To a lesser extent HCoV-OC43 also induced cell death signatures in responding cell subsets but no increase in these genes were observed following HCoV-NL63 infection. These results correlate with the H&E morphological analysis where both HCoV-229E and SARS-CoV-2/BA.5 induced the most tissue disruption. Increased expression of genes associated with several cell death pathways suggests that PANoptosis may be occurring in these cells.

### Pathogenic HCoV have distinct gene expression profiles compared to seasonal CoVs in edAT2 cells

To explore the host response to HCoV infection in edAT2 cells we performed bulk RNASeq analysis at 3 dpi. Given that the clinical HCoV-OC43 isolate did not replicate in edAT2 cells (Fig. 1F), we did not include HCoV-OC43 in this analysis. To remove any confounding effects of temperature on host response, this experiment was conducted at 35°C for both seasonal and pathogenic HCoVs. Some CPE was observed for MERS-CoV (∼50%) at this time point (data not shown). Like infectious virus titres in the supernatant, the level of viral genomes per sample were higher for the pathogenic HCoVs compared to the seasonal HCoVs (Fig. 7A). Principal component analysis (PCA) indicated that SARS-CoV-2 (B.1 and BA.5) and MERS-CoV clustered away from mock-inoculated and seasonal HCoV-infected edAT2 cells (Fig. 7B). MERS-CoV infection was distinguished by decreased expression of AT2 functional genes such as SFTPA1 and SFTPA2 that encode surfactant proteins and increased expression of genes associated with fibrosis such as KRT13 and KRT17 (Fig. 7C). In contrast, SARS-CoV-2 infection was associated with expression of innate immune genes such as NLRP9 and IFNAR (Fig. 7C). Pathway analysis reflected these differences in innate signalling (Sup. Fig 10A). Overall, these data suggest that pathogenic HCoVs may be driving differential gene expression relative to seasonal HCoVs in edAT2 cells.

Overall, all HCoVs influenced gene expression compared to mock-inoculated edAT2 cells with the seasonal HCoV inducing fewer significant changes (Sup. Fig. 10B). Of the significant differentially expressed genes, there were 75 significantly upregulated and 476 significantly downregulated by all five HCoVs (Fig. 7D). All viruses induced signalling pathways associated with induction and inhibition of antiviral responses and reduced the expression of genes associated with protein synthesis and oxidative phosphorylation confirming that HCoVs trigger innate signalling and hijack the host cell translational machinery for replication (Fig. 7E and Sup. Fig. 10C). Analysis of key interferon stimulated genes suggested there was a mixed response with pathogenic HCoVs suppressing OAS1 expression and all HCoVs increasing expression of TICAM1, ISG15 and IFIH1 (Fig.7H).

There were 1217 genes upregulated and 824 downregulated specifically by pathogenic HCoVs, whereas very few genes were modulated specifically by both HCoV-NL63 and HCoV-229E (Fig. 7D). Analysis of lung function pathways suggested that all viruses, except for HCoV-229E reduced the expression of genes associated with alveolar epithelial cell identity and signalling compared to mock (Fig. 7F). However, pathogenic HCoV had a more significant impact on AT2 identity with MERS-CoV also inducing gene expression indicative of fibrosis. Key surfactant genes such as SFTPA1 and SFTPB, were significantly downregulated following MERS-CoV infection (Fig. 7H). In contrast, genes associated with epithelial identity and injury such as EPCAM, CAPS and SMAD3 were modulated by all HCoVs, with trends towards greater responses in pathogenic HCoVs (Fig. 7H). Positive regulation of response to type II interferon was greater in pathogenic compared to seasonal HCoVs (Fig. 7G and Sup. Fig. 10D). Pathogenic HCoVs also significantly downregulated several inflammasome pathways suggestive of potential antagonism of the host immune response contributing to pathogenicity (Fig. 7G). Analysis of the genes associated with these pathways revealed that pathogenic HCoV infection led to reduced expression of CASP1, CASP5 and IL1B relative to mock and compared to HCoV-229E and HCoV-NL63 where no difference was observed (Fig. 7H). Another key difference between pathogenic and seasonal HCoVs was the extent of NF-ĸb signalling. SARS-CoV-2 and MERS-CoVs both significantly upregulated NFKB2, RELB, TNF, ICAM1 and CXCL8 compared to seasonal HCoVs (Fig. 7H). This suggests that pathogenic HCoVs induce enhanced NF-ĸB signalling compared to seasonal HCoVs and may reflect stronger inflammatory activation.

Interestingly, MERS-CoV shared a similar number of upregulated genes with HCoV-229E as it did with SARS-CoV-2 (Fig. 7D). Innate immune signalling signatures were higher in HCoV-229E and MERS-CoV infected edAT2 cells as well as genes associated with autophagy (Fig. 7G). FAS and TRAIL signalling were also strongly upregulated in MERS-CoV and HCoV-229E infected cells, correlating with the observed increase in LDH release and cytotoxicity following infection with these viruses (Fig. 2B). Furthermore, we found that both MERS-CoV and HCoV-229E strongly induced PKR/eIF2α, ATF4, PERK signalling pathways which are all key parts of the integrated stress response. Analysis of key cell death genes indicated that MERS-CoV induced expression of MLKL, FADD and FAS while HCoV-229E induced CASP8 and FADD expression (Fig.7H). This suggests that the type of cell death may be different between these two viruses.

Together these data show that pathogenic HCoVs caused greater disruption of AT2 cell function, particularly MERS-CoV, enhanced NF-ĸB signalling and distinct innate immune response differences compared to seasonal HCoVs. MERS-CoV and HCoV-229E also induced markers of enhanced cell death and stress responses suggesting that these two viruses may evoke similar host responses.

## Discussion

HCoVs can be broadly segregated into pathogenic and seasonal classifications based on their ability to cause severe respiratory disease in humans. While several factors likely drive the pathogenic potential of these viruses, the extent to which the response of the epithelium contributes to pathogenicity is unclear. Here, using specialised human airway epithelial cultures representing the nasal, bronchial, small airway and alveolar epithelium, we found that replication capacity and host responses to infection in the lower airways distinguishes pathogenic HCoVs from seasonal HCoVs. In addition, we identified several unique properties of HCoV-229E that distinguishes airway epithelial infection with this virus compared to other seasonal HCoVs.

Comparative analysis of virus replication capacity across the human respiratory tract showed that all HCoVs were able to replicate in human nasal epithelial cells. Consistent with a study by Otter *et al.,* we found that SARS-CoV-2 replicated to the highest titres in HNECs compared to MERS-CoV and seasonal HCoVs (*22*). In BCi and SABCi cultures, SARS-CoV-2 variants again replicated the most efficiently, with MERS-CoV restricted to replication in SABCi only, and all seasonal HCoVs exhibited comparatively restricted replication in the large and small airway epithelium. While MERS-CoV did not replicate in BCi cells, primary bronchial epithelial cells from three donors were permissive to MERS-CoV replication, suggesting that BCi cells may express insufficient DPP4 to permit infection (*29, 30*). Collectively, these data demonstrate that, beyond the universal ability of HCoVs to infect nasal epithelium, the capacity of HCoVs to replicate in proximal and distal airway epithelia varies markedly by virus and cell model.

CXCL10 and IFNs were the predominant cytokines produced following infection with HCoVs. CXCL10 is commonly expressed by airway epithelium in response to respiratory virus infection and can be used to predict virus infection in nasopharyngeal samples (*31, 32*). For SARS-CoV-2, CXCL10 expression was shown to play a critical role in disease resolution using a mouse model. In addition, serum CXCL10 levels can predict adverse outcomes in COVID-19 patients (*33, 34*). In our airway models, both pathogenic and seasonal HCoVs strongly induced CXCL10 expression, indicating that induction of this cytokine in the airway epithelium is not associated with pathogenicity. Cytokine expression was highest for all HCoVs in the nasal epithelium, with progressively reduced CXCL10 and IFN secretion by epithelial cells in response to infection in the large and small airways and lung. Low cytokine induction in edAT2 cells following HCoV infection is consistent with our previous studies (*21, 35*). These findings suggest that the lower lung epithelium may mount a less robust antiviral response, a phenomenon that has been previously demonstrated for rhinovirus infection (*36*). Consequently, it is conceivable that if viruses are able to reach and infect cells within the lower respiratory tract, they may encounter fewer epithelial-intrinsic barriers to replication and dissemination.

In this study we observed that pathogenic HCoVs produced higher levels of IFN-λ secretion compared to seasonal HCoVs in HNECs, BCi and SABCi cells. However, in our scRNAseq dataset, a drastic increase in responding cells with high mRNA levels of ISGs was observed following SARS-CoV-2, HCoV-229E and to a lesser extent HCoV-OC43. Therefore, despite evidence of lower IFN-λ secretion, HCoV-229E was able to robustly induce an ISG response. In contrast, MERS-CoV infection robustly suppressed ISG production despite detectable IFN-λ secretion. In addition, in SARS-CoV-2-infected cells there was a significant correlation between ISG expression and viral transcript levels suggesting that while this virus did not prevent ISG expression in responding cells, the virus could effectively shut down the IFN response in infected cells. This indicates a disconnect between IFN secretion and ISG expression in infected vs responding cells which was only decoupled through single cell analysis. In previous studies seasonal HCoVs were shown to induce robust IFN responses early during infection in primary nasal and bronchial cultures when compared to SARS-CoV-2 (*37*), which contrasts with our findings. In our study, we infected the airways models with pathogenic HCoVs at 37°C and seasonal HCoVs at 35°C to ensure optimal replication of each virus. Given that IFN signalling is reduced at lower temperatures (*38, 39*), this could partially explain the differences observed with our study. However, our results indicate that pathogenic HCoVs are better at shutting down the IFN response in infected cells than seasonal HCoVs even at 37°C.

Analysis of replication in edAT2 cells distinguished the pathogenic HCoVs from the seasonal HCoVs, with SARS-CoV-2 and MERS-CoV showing high viral titres compared to low peak titres for HCoV-229E and HCoV-NL63 and no replication for HCoV-OC43. These data align with previous reports that SARS-CoV, MERS-CoV and SARS-CoV-2 can efficiently infect human alveolar tissue ex vivo and AT2-like cultures (*40–45*), whereas seasonal HCoVs are largely confined to the upper airways. While all HCoVs induced NF-κB signalling and suppressed protein synthesis and oxidative phosphorylation in edAT2 cells, pathogenic HCoVs induced a more robust NF-κB response and changes to AT2 function and signalling. They also enhanced type II interferon responses while suppressing inflammasome-associated genes, including CASP1, CASP5 and IL1B, suggesting that differential modulation of innate immunity may contribute to pathogenicity. MERS-CoV showed the most distinct response, with extensive remodelling of AT2 identity and function, fibrosis, cytoskeletal remodelling and apoptosis. MERS-CoV and HCoV-229E both strongly induced FAS, TRAIL and TNF signalling, as well as PKR/eIF2α–ATF4–PERK, unfolded protein and DNA-damage responses, consistent with substantial cellular stress and cell death. Thus, while cell death is a shared feature of some HCoVs, pathogenicity likely reflects the combined effects of epithelial dysfunction, immune modulation, metabolic disruption and cell death rather than cell death alone.

Among the seasonal HCoVs, HCoV-229E displayed several unexpected features that distinguished it from other seasonal and pathogenic HCoVs. Despite only modest replication in airway epithelial cultures and lower peak titres in edAT2 cells than SARS-CoV-2 and MERS-CoV, HCoV-229E caused disproportionately high levels of epithelial damage, with the greatest LDH release across the airways and marked cytopathic effects in edAT2 cells. In HNECs, HCoV-229E infection led to extensive loss of cilia and squamous epithelial morphology, consistent with reduced tissue integrity and dedifferentiation (*46*). ScRNAseq showed that HCoV-229E-infected ciliated cells down-regulated key ciliated cell identity genes and up-regulated genes associated with antiviral and stress pathways, and infected cells accumulated in G1 rather than S phase. HCoV-229E also showed robust activation of cell death pathways in both nasal epithelium and edAT2 cells, closely resembling MERS-CoV in its ability to trigger apoptosis and stress responses despite its poorer replication. This suggests that while HCoV-229E has a profound effect on cellular function compared to other seasonal HCoVs, this still does not increase replication capacity. There is limited evidence that HCoV-229E is more virulent in humans compared to other seasonal HCoVs. In a small cohort of 133 patients, HCoV-229E was associated with an approximately 3-fold increased risk of mortality compared to HCoV-OC43 (*47*). Collectively, our findings suggest that HCoV-229E has unique virus-host interactions in the human respiratory tract compared to other seasonal HCoVs, combining limited alveolar replication with a propensity to disrupt epithelial cell identity and cell survival.

Unlike conventional immortalized cell lines, airway culture models more closely recapitulate the cellular composition, architecture, differentiation state, and functional characteristics of the human airway epithelium. Consequently, they provide a physiologically representative platform for investigating virus-host interactions, including viral entry, replication, innate immune activation, and host-mediated antiviral responses. However, despite the advantages of these systems, the sample size of primary human airway cultures used in this study are limited (n=2-3 for nasal and bronchial cultures). Moreover, scRNAseq was only performed on one HNEC donor. In addition, the BCi, SABCi and edAT2 systems are highly reproducible culture systems but represent one human donor. Therefore, the results generated using these culture systems are not representative of the entire human population. We also noted that the proportion of infected cells were overall low in HNECs particularly for HCoV-NL63 and HCoV-OC43 despite detectable virus release making it difficult to analyse infected populations across all the virus strains. Similarly, despite consistent levels of viral transcripts the proportion of infected vs bystander cells within the edAT2 bulk RNASeq data may also differ between virus strains. Moreover, these systems are purely epithelial cell culture systems and lack immune cells, fibroblasts and endothelial cells. Future studies should incorporate these non-epithelial cell types to understand how different HCoVs influence cellular cross-talk and how this influences responses to infection. Another critical factor when comparing HCoVs relates to assumption that one virus isolate is representative of all isolates from that viral family or that lab-adapted strains will respond similarly to contemporary strains. All of these aspects should be considered when using these models to study respiratory virus infection.

In summary, our comparative analysis across physiologically relevant human airway models demonstrates that the replication to high titres in AT2 cells, together with a distinct pattern of epithelial responses can segregate pathogenic from seasonal HCoVs. In addition, we show that pathogenicity is not explained solely by cytotoxicity, since HCoV-229E, despite modest replication, induced profound epithelial damage, ciliary loss and stress/death programmes akin to MERS-CoV. Our findings underscore the importance of assessing viral infection and replication kinetics in distal lung epithelium and integrating epithelial host responses with immune and stromal interactions, to refine risk assessment, guide therapeutic prioritisation and improve pandemic preparedness for emergent coronaviruses.

## Materials and Methods

### Cell lines

African green monkey kidney epithelial (Vero) cells (American Type Culture Collection (ATCC), Cat. CCL-81) were maintained in Minimum Eagle Medium (MEM, Media Preparation Unit, Peter Doherty Institute) supplemented with 5% (vol/vol) Fetal Bovine Serum (FBS, Corning), 50U/ml Penicillin and 50µg/ml Streptomycin (Gibco), 2mM GlutaMAX (Gibco) and 15 mM HEPES (Gibco). Vero cells expressing human signalling lymphocytic activation molecule (VerohSLAM, European Collection of Authenticated Cell Cultures (ECACC), Cat. 04091501) were maintained in Minimum Eagle Medium supplemented with 7% (vol/vol) FBS, 50U/ml Penicillin and 50µg/ml Streptomycin, 2mM GlutaMAX, 15 mM HEPES and 0.25mg/mL Geneticin (Gibco). VeroE6 cells expressing transmembrane serine protease TMPRSS2 (VeroE6-TMPRSS2 cells) were maintained in Dulbecco’s Modified Eagle Medium (DMEM, Media Preparation Unit, Peter Doherty Institute) supplemented with 10% (vol/vol) FBS, 50U/ml Penicillin and 50µg/ml Streptomycin, 2mM GlutaMAX, and 1mg/mL Geneticin. LLC-MK2 cells were maintained in MEM supplemented with 50 U/mL penicillin, 50 µg/mL streptomycin, 15 mM HEPES, 2 mM Glutamax, and 5% (vol/vol) FBS. Human fetal lung fibroblast cells (MRC5) were maintained in DMEM supplemented with 10% (vol/vol) FBS, 50U/ml Penicillin and 50µg/ml Streptomycin, 2mM GlutaMAX, 0.1mM Non-Essential Amino Acids (Gibco) and 1mM Sodium Pyruvate (Fisher Scientific). Human hepatocyte-derived carcinoma cells (Huh7) were maintained in DMEM supplemented with 10% (vol/vol) FBS, 50U/ml Penicillin and 50µg/ml Streptomycin and 2mM GlutaMAX. Human lung cancer cells (Calu3) were maintained at 37°C, 5% CO2 in Eagle Minimum Essential Medium (EMEM, Gibco, Cat. M4655) supplemented with 10% (vol/vol) FBS, 50U/ml Penicillin and 50µg/ml Streptomycin, 0.2mM non-essential amino acids and 0.5mM Sodium Pyruvate. All immortalised cells were maintained at 37°C with 5% CO2.

### Primary nasal epithelial cells

Healthy adults were recruited for this study under ethics approval from the Royal Children’s Hospital (RCH) human ethics committee (HREC/35132). Nasal epithelial cells were sampled and cultured as described previously (*27, 48, 49*).

### Primary bronchial epithelial cells

Bronchial airway brushings were collected from patients undergoing routine diagnostic bronchoscopy for suspected lung cancer at the Royal Melbourne Hospital respiratory clinic as previously described (*50*). Ethics approval was obtained from the Melbourne Health Human Research Ethics Committee (2020.112) and the School of Biomedical Sciences Human Ethics Advisory Group (HEAG; 1851394) at the University of Melbourne. Briefly, a sheathed cytology brush was gently wedged in a peripheral bronchus and withdrawn 2 to 3 cm before the brush tip was advanced 2 to 3 cm beyond the protective sheath. Gentle brushing was performed, after which the brush tip was retracted back into the sheath, and the device was fully removed. The brush tip was detached into DMEM (Gibco) and the procedure was repeated as required. Cells were pelleted by centrifugation, resuspended in PneumaCult Ex Plus Medium (Stemcell Technologies) supplemented with 96 ng/mL Hydrocortisone (StemCell Technologies) and 50U/ml Penicillin and 50µg/ml Streptomycin, and seeded into cell culture flasks coated with 2 μg/mL of human placental collagen (Sigma-Aldrich).

### Immortalised airway epithelial cells

Basal-like human airway progenitor cells (BCi-NS1.1 (*51*)) and human small airway basal cells (hSABCi-NS1.1 (*52*)), a kind gift from Professor Ron Crystal, Cornell University, NY, were maintained at 37°C, 5% CO2 in Pneumacult-Ex Plus medium and used to generated human airway epithelial cultures. To differentiate human airway epithelial cultures, BCi and hSABCi cells were seeded at 75,000 cells/well in transwell membrane supports (6.5 mm, Corning) and cultured in Pneumacult-Ex Plus medium in apical and basolateral chambers. Upon reaching confluence (2-3 days post-seeding), apical medium was removed to expose cells to ambient air; medium in the basolateral chamber was replaced with Pneumacult ALI maintenance medium (Stemcell Technologies) supplemented with 4µg/mL Heparin (Stemcell Technologies), 480 ng/mL Hydrocortisone and 50U/ml Penicillin and 50µg/ml Streptomycin. Medium in the basolateral chamber was replaced with fresh Pneumacult ALI maintenance medium every 2 to 3 days. Cells were cultured for a minimum of 28 days to form differentiated, polarized cultures that mimic in vivo pseudostratified mucociliary epithelium. The apical surfaces of cultures were washed to remove mucus with PBS (+Ca2+/Mg2+) at 14-and 21-days post-airlift. BCi and hSABCi cultures were used for SARS-CoV-2 infection following 4 weeks of differentiation at an air-liquid interface (ALI).

### Generation of hESC-derived AT2 cells

Human embryonic H9 (female) stem cells were differentiated into edAT2 cells as previously described (*27, 35*). Briefly, H9 stem cells were differentiated as a three-dimensional organoid for 18 days in flasks coated with Matrigel (Corning). Lung organoids were maintained for experiments between passage 2-8 and dissociated with TrypLE (Thermo Fisher Scientific) and seeded onto Geltrex-coated plates (Gibco). Cells were then maintained for a further 7-10 days in two-dimensional culture until infection at >70% confluency.

### Viruses

SARS-CoV-2 B.1 (hCoV-19/Australia/VIC01/2020, GISAID: EPI_ISL_406844) stocks was grown in VerohSLAM cells at 37°C in serum-free MEM (containing 50U/ml Penicillin and 50µg/ml Streptomycin and 2mM GlutaMAX) in the presence of 1µg/ml TPCK-Trypsin (Worthington Biochem). SARS-CoV-2 BA.5 (hCoV-19/Australia/VIC61194/2021, GISAID: EPI_ISL_13276063) was grown in Calu3 cells at 37°C in infection media (2% FBS, 0.2mM Non-Essential Amino Acids, 50U/ml Penicillin and 50µg/ml Streptomycin, 0.5mM Sodium Pyruvate). MERS-CoV was grown in Vero cells at 37°C in serum-free MEM (containing 50U/ml Penicillin and 50µg/ml Streptomycin and 2mM GlutaMAX) in the presence of 1µg/ml TPCK-Trypsin. HCoV-NL63-Am1 stocks were grown in LLC-MK2 cells at 35°C in MEM supplemented with 2% FBS, 50U/ml Penicillin and 50µg/ml Streptomycin, 15mM HEPES and 2mM GlutaMAX. HCoV-229E (VR-740) was prepared from supernatants of infected MRC-5 cells in MEM supplemented with 1% FBS, 50U/ml Penicillin and 50µg/ml Streptomycin, 0.1mM non-essential amino acids and 1mM Sodium Pyruvate and 2mM GlutaMAX. A clinical HCoV-229E isolate (229E/Australia/22050721/2022) was propagated in Huh7 cells supplemented with 1% FBS, 50U/ml Penicillin and 50µg/ml Streptomycin, 2mM GlutaMAX and 1 µg/mL TPCK-trypsin. Virus stocks were harvested at 70-90% cytopathic effect (CPE) in cultures, centrifuged at 2000rpm for 5 minutes to remove cellular debris, aliquoted into cryovials and stored at-80°C. A clinical HCoV-OC43 isolate (OC43/Australia/19052914/2019) was isolated using BCi Cell as before(*21*). Stocks were titred as described below. All infectious virus work with SARS-CoV-2 and MERS-CoV was performed inside a biosafety II cabinet in a biosafety containment level 3 facility with approval from the institutional biosafety committee. Personnel wore powered air-purifying respirators (3M TR-315A VERSAFLO) or P2 masks.

### Virus Titrations

SARS-CoV-2 titres were determined using an infectivity assay in VeroE6-TMPRSS2 cell monolayers with serum-free DMEM containing 1µg/ml TPCK-Trypsin. Cells were incubated at 37°C and assessed for CPE on day 4. MERS-CoV titres were determined using an infectivity assay in Vero cell monolayers with serum-free MEM containing 1µg/ml TPCK-Trypsin. Cells were incubated at 37°C and assessed for CPE on day 4. HCoV-229e titres were determined using an infectivity assay in MRC-5 cell monolayers with DMEM containing 1% FBS and 1µg/ml TPCK-Trypsin. Cells were incubated at 35°C and assessed for CPE on day 4. HCoV-NL63 titres were determined using an infectivity assay in LLC-MK2 cell monolayers with MEM containing 2% FBS and 1μg/mL TPCK-trypsin. Cells were incubated at 35°C and assessed for CPE on day 7. Virus titres are expressed as mean log10 TCID50/ml.

### RT-qPCR for clinical OC43

RNA was extracted as per the manufacturer’s recommendation using Qiagen QiaCube HT and QiaAmp 96 Virus QiaCube HT kit. RT-PCR reaction was setup using SensiFast Probe No-ROX One-Step Kit (Bioline) using the following primers/probes: OC43 Forward: 5’-TGTTTATGGCTGATGCTTATCT-3’, OC43 Reverse: 5’-AAGGTATTACACATACCGCAAA-3’ and OC43 Probe: 5’-CACTGTGTGGTATGTGGGGCAAAT-3’. Serial 10-fold dilutions of plasmid encoding the OC43 E gene were used to generate a standard curve and calculate the virus genome copies in the samples.

### Virus infection

Prior to infection, airway cultures were washed once with pre-warmed PBS (+Ca2+/Mg2+) to remove mucus on the apical surface. Cells were inoculated with 100µl virus diluted to a multiplicity of infection (MOI) of 0.1 for 2 hours at 37°C for SARS-CoV-2 and MERS-CoV and 35°C for HCoV-NL63 and HCoV-229E. For HCoV-OC43, cells were inoculated with 106 genome copies per well for 2 hours at 35°C. The inoculum was removed, and the apical surface washed three times with PBS and the third wash collected as the day 0 sample. On days 1-7 post-infection, 200µl PBS was added to the apical chamber and incubated for 10 minutes before samples were stored at-80°C. Basolateral medium was replaced every second day with ALI maintenance medium. Apical samples were assayed for infectious virus by virus titration or RT-qPCR. For infection of edAT2 cells, cells were inoculated with 10^4^ TCID50 of virus in 100µl for 1 hour at 37°C for SARS-CoV-2 and MERS-CoV and 35°C for HCoV-NL63 and HCoV-229E. For HCoV-OC43, cells were inoculated with 106 genome copies per well for 1 hour at 35°C. Cells were washed three times with PBS, the third wash was collected as the day 0 sample and cells were re-fed with lung edAT2 medium. Supernatant samples were collected every day for seven days and media was replenished on days 1-5. Supernatant samples were assayed for infectious virus by virus titration as described above and replication by qPCR.

### Cytokine secretion and cell death analysis

Cytokine/chemokine concentrations in supernatants or basolateral medium were analysed using the LEGENDplex human anti-virus response panel (Biolegend) following the manufacturer’s instructions. Samples were run on a BD FACSCanto II and analysed using LEGENDplex™ Data Analysis Software Suite. LDH was measured in supernatants or basolateral medium using the LDH cytotoxicity kit (Abcam) according to the manufacturer’s instructions using an LDH standard curve.

### Immunohistochemistry

Airway epithelial cultures in transwells were fixed in 10% formalin for 30 minutes at room temperature before three PBS washes were performed. Membranes were cut from transwells, embedded in paraffin and 5µm cross-sections were cut using a Microm HM 325 paraffin microtome onto Menzel Gläser, SuperFrost® Plus glass slides (Rowe Scientific). Sections were deparafinised and rehydrated by immersing in xylene twice for 10 minutes each, 100% ethanol twice for 10 minutes each followed by 95% ethanol, 70% ethanol and 50% ethanol for five minutes each. Slides were washed in PBS twice and stained for hematoxylin and eosin. To retrieve antigens, slides were submerged in Universal HIER antigen retrieval buffer (Abcam) and covered. Slides were heated in a microwave (100% power for 3 minutes, 90% power for 2 minutes) and incubated for 9 minutes covered before being cooled on a shaker and rinsed three times with deionized water. For detection of antigens using immunohistochemistry, slides were blocked with 5% normal goat serum (Thermo Fisher Scientific) containing 0.1% Triton X-100 (Sigma Aldrich) in PBS for 1 hour at room temperature. Slides were washed three times in PBS and stained with primary antibodies overnight at 4°C. Primary antibodies used were α-acetylated tubulin (Sigma Aldrich, 1:1000), α-Mucin 5AC (Abcam, 1:500), α-Club cell protein 10 (R&D Systems, 1:100), α-SARS-CoV Nucleocapsid (Sino-Biological, 1:500), α-MERS-CoV Nucleocapsid (Sino-Biological, 1:500), α-HCoV-229E Nucleocapsid (Sino-Biological, 1:500), α-HCoV-NL63 Nucleocapsid (Sino-Biological, 1:500) and α-HCoV-OC43 Nucleocapsid (Sino-Biological, 1:500). Slides were washed three times in PBS and incubated with secondary antibodies at room temperature for 1 hour in the dark. Secondary antibodies used were acquired from Thermo Fisher Scientific: Goat anti-Rabbit IgG (H+L) AF568 (1:500), Goat anti-Rat IgG (H+L) AF488 (1:500), Goat anti-Mouse IgG1 (H+L) AF647 (1:500) and Goat anti-Mouse IgG2b (H+L) AF488 (1:500). Slides were washed three times with PBS and counter stained with 4’,6-Diamidino-2-Phenylindole, Dihydrochloride (DAPI, Thermo Fisher Scientific) for 5 minutes at room temperature in the dark. Slides were washed three times with PBS and mounted with glass cover slips using Vectashield mounting medium (Abacus). Slides were imaged on a Zeiss LSM780 or LSM980 confocal microscope and analysed with Zen software and ImageJ.

### Single cell RNA sequencing

Human nasal epithelial cells, BCi and SABCi cells differentiated at an ALI were inoculated as described above. At 1- and 3-dpi, cells were washed once with warm PBS for 10 minutes to harvest apical wash samples followed by two apical washes with warm Hanks Balanced Salt Solution (HBSS, Doherty Institute Media Preparation Unit) for 5 minutes each to remove extracellular mucus. The apical and basolateral chambers were washed with HBSS before incubation with Animal Component-Free Dissociation Reagent (Stemcell Technologies) for 10 minutes. Dissociated cells were transferred to a 15ml Tube containing Animal Component-Free Inhibition Reagent (Stemcell Technologies). A total of three rounds of dissociation were performed to completely remove cells from the transwell insert. Cells were washed three times with ALI Maintenance medium containing 0.1% Pluronic (ThermoFisher). 500,000 cells from each condition were incubated with TotalSeq™-C0251 anti-human Hashtag antibodies 1-6 (BioLegend, 1:500 dilution) for 30 minutes on ice. Cells were washed twice with PBS containing 10% FBS and once with PBS containing 0.01% BSA. Cells were pooled (6 samples per pool), loaded to the 10X Genomics Chromium Controller (approximately 10,000 cells loaded to 1 Chromium lane) and processed with the Chromium Next GEM single-cell 5′ library and gel bead kit according to manufacturer’s protocols. Gene expression and Cell Surface Protein libraries were pooled and sequenced in multiplex on the Illumina NextSeq 2000 system with the following read configuration: Read 1: 26 cycles, i7 Index: 10 cycles, i5 Index: 10 cycles and Read 2: 90 cycles. Gene expression libraries were sequenced to an estimated depth of 25,000 read pairs per cell, and Cell Surface Protein libraries were sequenced to an estimated depth of 5,000 read pairs per cell.

## Single Cell RNA Sequencing Analysis

### Read mapping and quantification

Raw base call files were converted to FASTQ format and demultiplexed using the Cell Ranger ‘mkfastq’ pipeline. To ensure consistent cell identification across modalities, FASTQ files from gene expression and feature barcode libraries were integrated using the Cell Ranger ‘multi’ command. Gene expression reads were aligned to the GRCh38 human reference genome (10x Genomics) for single-cell gene expression (scRNA-seq) analysis. The integration of libraries was done by the Cell Ranger ‘merge’ command, which can normalise the library depth difference. The quantification of viral infection was performed by mapping the gene expression libraries to corresponding references: B.1 (MT007544.1: Severe acute respiratory syndrome coronavirus 2 isolate SARS-CoV-2/human/AUS/VIC01/2020), BA.5 (OP165284.1: SARS-CoV-2/human/USA/IN-CDC-STM-WR9QFA336/2022), HCoV-NL63 (NC_005831.2: Human Coronavirus NL63), OC43 (LC756670.1: HCoV-OC43_Fukushima_OR427_2021), HCoV-229E (ON791801.1: Human coronavirus 229E strain LC22), and MERS-CoV (NC_019843.3: Middle East respiratory syndrome-related coronavirus isolate HCoV-EMC/2012). The resulting BAM files aligned to the human reference were analysed for single-nucleotide polymorphism (SNP) patterns to verify sample identity using the ‘vireoSNP’ package.

### Data preprocessing and integration

Count matrices were processed using Seurat to filter out low-quality cells and those with high mitochondrial content. We utilised ‘MULTISeqDemux’ for feature barcode demultiplexing, integrating library and oligonucleotide identities to define experimental conditions. Assignments were validated against genotype demultiplexing, with doublets and negative cells excluded from the study. For data integration, we used rliger to perform non-negative matrix factorisation (k=30, lambda=5, convergence threshold= 10e-6), allowing for the identification of shared and dataset-specific factors. To visualise the integrated data, Uniform Manifold Approximation and Projection (UMAP) was performed on the aligned cell factor loadings. The final object was subset to retain only HNEC cell systems for downstream analysis.

### Data clustering and annotation

Unsupervised clustering was performed on the integrated factor loadings using the Leiden algorithm at a resolution of 0.8. Batch identities were overlaid on the resulting clusters to ensure that the grouping was not driven by batch effects. Cell type markers were determined by identifying the top five genes associated with each loading factor and validated through literature review and transcript profiles.

### Differential gene expression analyses

Differentially expressed gene analysis was performed to compare the exposed cells to mock controls and infected cells to non-infected cells using a pseudo-bulk approach via ‘rliger’ framework with the n pseudo-replicates of 30 to avoid the pitfall when there is no replication in some samples. Significance was defined by a Benjamini-Hochberg adjusted p-value (p_adj < 0.05) and an absolute log2 fold-change threshold |log_2FC| > 2). Gene ontology (GO) and pathway annotations were investigated using enrichGO and enrichKEGG from the clusterProfiler R package.

### Gene set enrichment analyses

Single-sample Gene Set Enrichment Analysis (ssGSEA) was utilized to quantify functional states at single-cell resolution. Using the escape R package, enrichment scores were calculated for the HALLMARK_INTERFERON_ALPHA_RESPONSE and HALLMARK_INTERFERON_GAMMA_RESPONSE pathways. To ensure consistency across experimental batches, scores were calculated within individual batches using normalized data before being integrated. To distinguish between inflammatory states, an interferon-response threshold was established at the 95th percentile of the mock control enrichment scores. Cells exceeding this threshold were classified as IFN-responders, while those below were designated as non-responders.

### RNA velocity analyses

Spliced and unspliced transcript matrices were generated from individual BAM files using velocyto and integrated with curated metadata from the previous section. Using scVelo, we filtered genes with a minimum of 20 shared counts and calculated first-and second-order moments based on a 30-neighbor graph. Batch-Balanced k-Nearest Neighbours was used for batch correction (*53*). Transcriptional kinetics were resolved using the stochastic model to estimate gene-specific kinetic rates and project velocity vectors onto the UMAP embedding. To resolve the cluster-level lineage relation, the neighbours from K-nearest Neighbours graph were used to compute the most frequent neighbour cell types.

### Bulk RNA Sequencing in edAT2 cells

edAT2 cells were infected as described above at 35°C. At 3 dpi, each well was harvested into 500µL of TRI reagent and RNA extracted using the Direct-zol RNA Miniprep (Zymo Research). Samples (3 replicates per condition) were sequenced using the Illumina NextSeq 2000 system to an estimated depth of 20M reads per sample.

## Bulk RNA Sequencing Analysis

### Pre-processing RNA-seq

Raw next-generation paired-end RNA-seq reads were obtained in the FASTQ format and were quality trimmed with with fastp v.13.6 using default parameters for paired-end reads (*54*). Following this, trimmed reads were aligned with STAR v2.7.11b to a combined reference (*55*), which is comprised of the human genome (GRCh38, Ensembl release 110 primary assembly) and the genomes of MERS-CoV (EMC/2012), SARS-CoV-2 B.1 (ancestral, Australia/VIC01/2020), SARS-CoV-2 Omicron BA.5 (Australia/VIC61194/2022), HCoV-229E (229E/Australia/22050721/2022) and HCoV-NL63 (Amsterdam-1/2004), with the local alignment parameters (--alignEndsType Local), and maximum intron size 1Mb. Alignments were then sorted, and duplicates were markerd and filtered with samtools v1.23.1 (*56*) with parameters fixmate-m, markdup-r (*57*). Read counts were then obtained with Subread’s v2.1.1 featureCounts in paired-end mode (*58*). Quality control was summarised with MultiQC v1.35 (*59*).

## Bulk RNA-seq gene expression analyses

### Counts processing and normalisation

The gene level counts were analysed in R v4.6.1. Reads assigned to the viral genes were removed from the count matrix prior to processing. Ensembl gene IDs were annotated with HGNC gene symbols using biomRt v2.68.0 (*60*). Lowly expressed genes were removed with edgeR’s v4.10.1 (*61–63*) filterByExpr, using the experimental group as a grouping factor. Libraries were then trimmed mean of M-values (TMM) normalised (*64*). Normalised expression was then expressed as log2 counts per million (log2[CPM]).

### Viral transcript quantification

Viral read counts were normalised separately, using the TMM-adjusted effective library sizes of the host genes. They were then expressed as log2[CPM], and not utilised in further analyses.

### Dimension reduction

Principal component analysis (PCA) was performed on the log2[CPM] values with prcomp (base R). Correspondence analysis (COA) was performed with made4 v.1.86.0 (*65, 66*). For each of the first two COA axes (C1, C2), the top 500 genes with the most negative, and 500 genes with the most positive coordinates were tested for Gene Ontology (GO) Biological processes (BP) enrichment (*67*) using topGO v2.64.0 (*68*), with the weight01 algorithm, and genes were mapped to GO terms with the Bioconductor package org.Hs.eg.db.

### Gene set variation analysis (GSVA)

For pathways of interest, GSVA scores were calculated for each condition with GSVA v2.6.6 (*69*), using group mean log2[CPM]. Gene sets were obtained from the Molecular Signatures Database (MSigDB with mdigdbr v26.1.1 (*70*). Lung function was assessed with AT2 cell signatures from the Human Lung Cell Atlas (*71*), as well as selected GO terms and WikiPathways (*72*) related to lung function and fibrosis. Immune activity was assessed with curated immune associated gene sets from GO BP, Reactome (*73*), the Pathway Interaction Database (*74*), KEGG MEDICUS (*75*) and Wikipathways. The top ten most variable gene sets across samples were selected from each collection for presentation.

### Differential Gene Expression (DGE) analyses

DGE analyses were performed with edgeR’s quasi-likelihood framework (*76*), with a no intercept design and one term per condition. Contrasts were defined with limma v3.68.4 (*77*). P-values were adjusted with the Benhamini-Hochberg method. Genes with a false discovery rate (FDR) <0.05 were considered differentially expressed.

### Gene set enrichment analysis (GSEA)

For each HCoV versus Mock DGE comparison, genes were ranked by sign(log2(fold change)) *-log10(FDR), Pre-ranked GSEA was then performed with fgsea v1.38.0 (*78*), using GO BP and Reactome gene sets. Common pathways were defined as gene sets tested in all fie comparisons as significant (adjusted P < 0.05) in at least three. The top 5 terms with the highest, and top five with the lowest mean normalised enrichment scores (NES) were reported.

### Visualisation

Figures were generated with ggplot2 v4.0.3, pheatmap v1.0.13, ComplexHeatmap v.2.28.0 (*79*) and venn v.1.13.

## Statistical Analysis

Data was analysed using GraphPad Prism Version 10. Virus replication over time was analysed using Area Under Curve analysis followed by one-way ANOVA with Dunnett’s multiple comparisons test. Cytokine expression analysis was performed using a student t-test. P values are shown either in the figure or described in the figure legends.

## Acknowledgements

SARS-CoV-2 B.1 and BA.5 variants were kindly provided by the Victorian Infectious Diseases Reference Laboratory (VIDRL). MERS-CoV was a kind gift from Prof Malik Pieris, University of Hong Kong. HCoV-NL63-Am1 (Amsterdam-1) was a kind gift from Prof Lia van der Hoek, University of Amsterdam. HCoV-229E (VR-740) and HCoV-OC43 (VR-1558) were kind gifts from Prof Nathan Bartlett, University of Newcastle. This work was supported by National Health and Medical Research Council of Australia (Investigator fellowship, KS), the Cumming Global Centre for Pandemic Therapeutics (JAN and KS) and a Doherty Institute Collaborative Seed grant (MJG, SS, AF and JAN). The Melbourne WHO Collaborating Centre for Reference and Research on Influenza is supported by the Australian Centre for Disease Control.

