## Supplementary material for "Viral replication and host response in airway epithelial cells predicts coronavirus pathogenicity and identifies HCoV-229E as an atypical seasonal coronavirus": Supplmentary Files

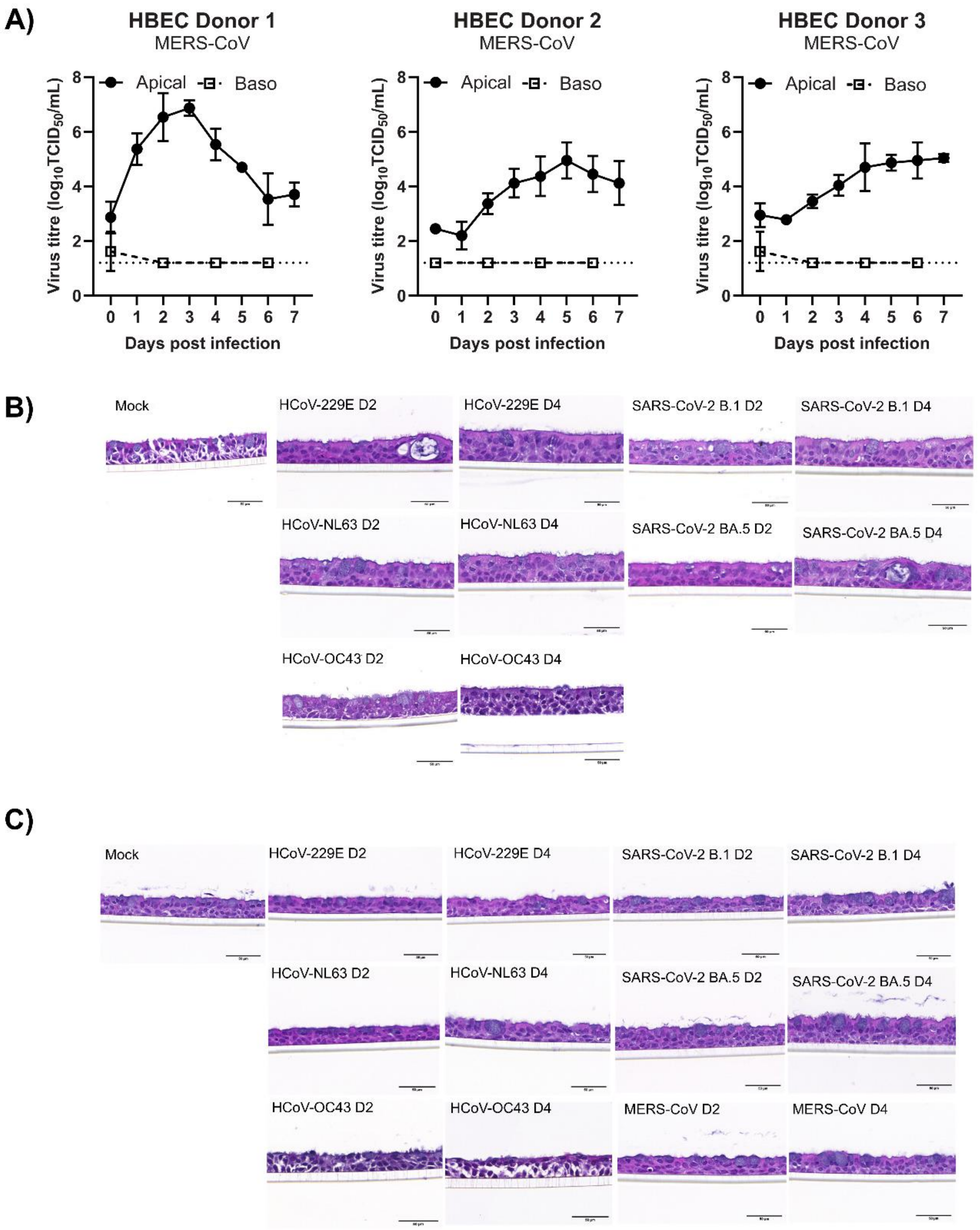


**Supplementary Figure 1: MERS-CoV replicates robustly in primary human bronchial cells. (A)** MERS-CoV replication following infection of three primary human bronchial epithelial cell donors shown as $\log_{10}$ 50% tissue culture infectious units per ml ($TCID_{50}$/ml). Solid circles represent titres in apical wash samples and open squares represent titres in basal medium. Mean ± SD is shown from three technical replicates. Dotted lines represent lower limit of detection. Representative images of BCi **(B)** and SABCi **(C)** sections infected with HCoV-229E, HCoV-NL63, HCoV-OC43, SARS-CoV-2/B.1, SARS-CoV-2/BA.5 and MERS-CoV fixed at 2 (D2) or 4 (D4) dpi and stained with haematoxylin and eosin. Scale bar = 50 μm.

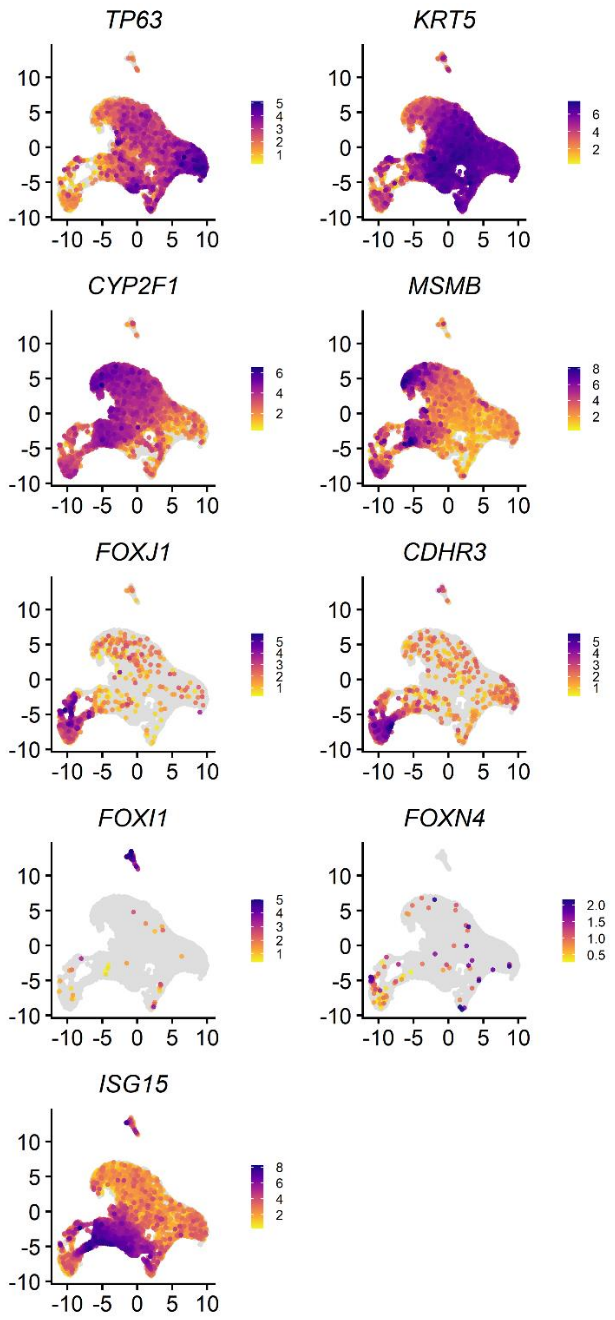


**Supplementary Figure 2: Expression of genes associated with different epithelial cell types.** Uniform Manifold Approximation and Projection (UMAP) of all cells profiled by single-cell RNA sequencing. Each dot represents one cell, coloured by normalised expression of: basal epithelial markers (TP63 and KRT5), cycling basal cell marker MKI67, club cell markers (SCGB1A1 and CYP2F1), goblet cell markers (MSMB, TFF3 and MUC5AC) ciliated cell markers (CDHR3, FOXJ1 and DNAH9), ionocyte markers (CFTR and FOXI1), deuterosomal cell markers (FOXN4 and DEUP1), ER stress marker DDIT3 and the antiviral gene ISG15.

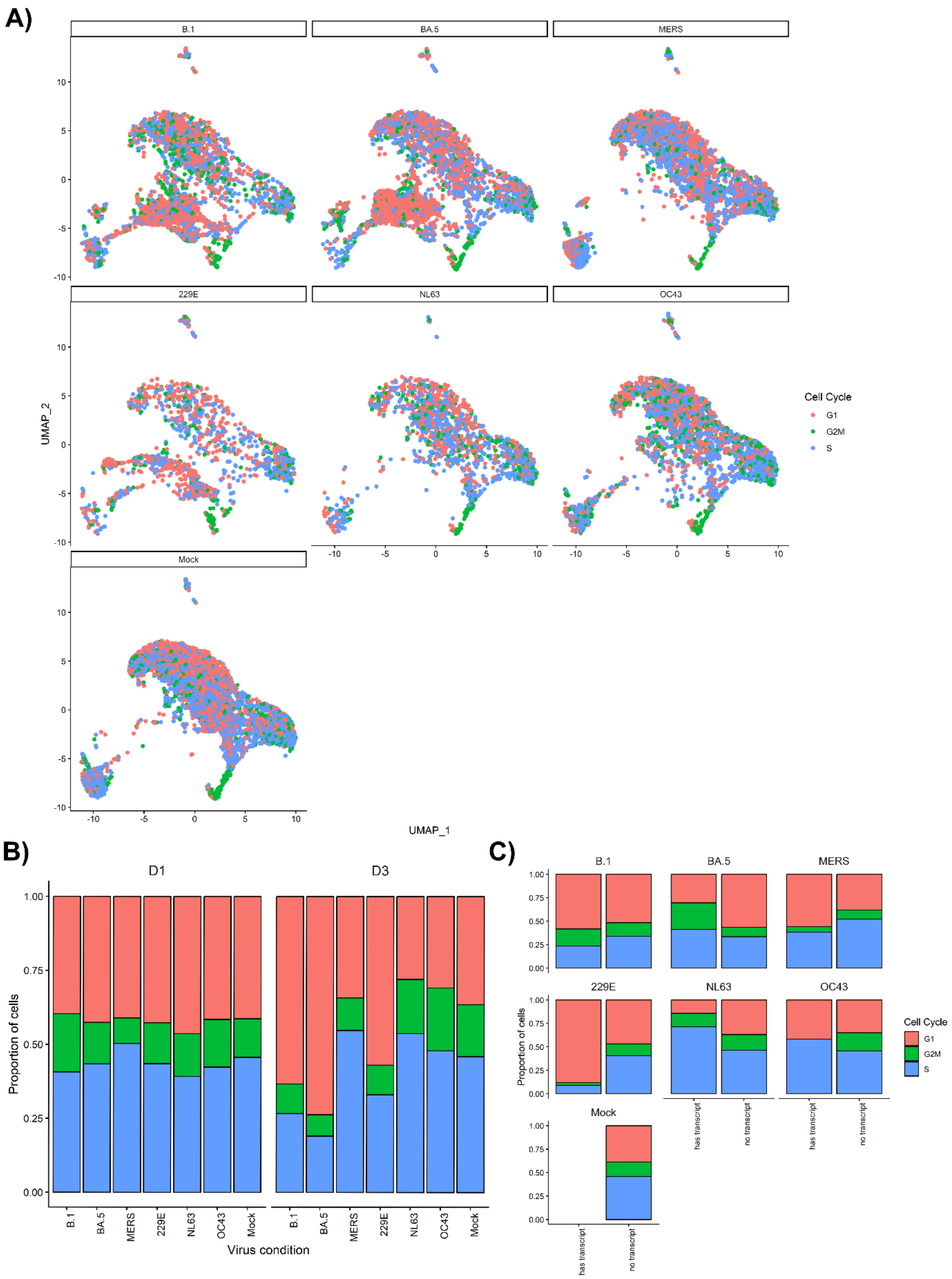


**Supplementary Figure 3: SARS-CoV-2 and HCoV-229E increase the proportion of cells in G1 phase. (A)** UMAP representation of cells from each experimental condition. Red dots represent cells in G1 phase; green dots represent cells in G2M phase and blue dots represent cells in S phase. Bar plots of the relative proportion of cells in each cell cycle phase across time points in all cells **(B)** or only virus transcript-containing cells **(C)** from each condition. For **(B),** the x axis represents the different viruses, and the y axis indicates the proportion of each cell cycle phase, with the total proportions summing to 100% per virus on 1 and 3 dpi. For **(C)**, the x axis represents presence or absence of viral transcripts, and the y axis indicates the proportion of each cell cycle phase, with the total proportions summing to 100% per virus on 3 dpi.

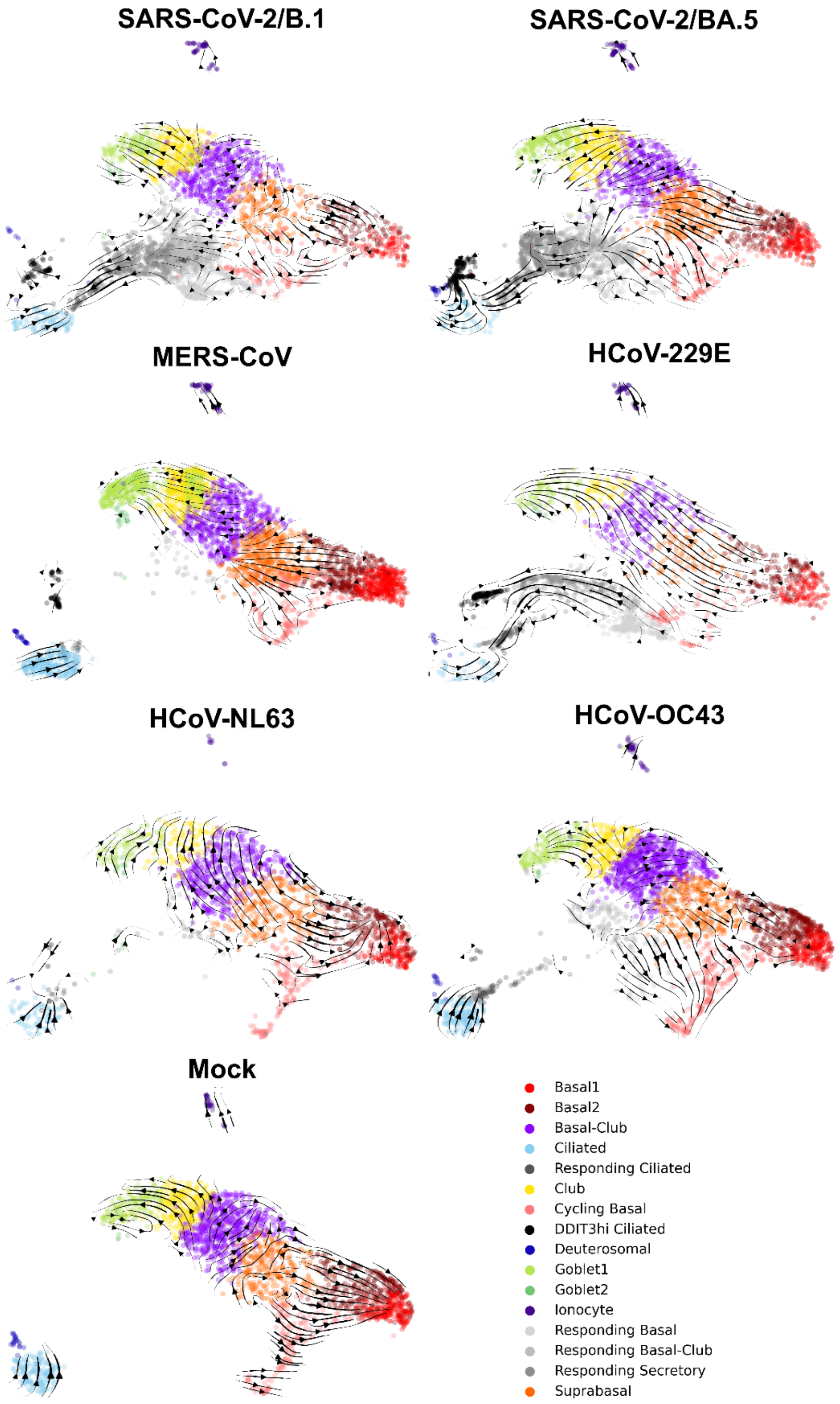


**Supplementary Figure 4: Cellular differentiation trajectories inferred by RNA Velocity.** RNA velocity estimation for all epithelial cell types split by cells inoculated with SARS-CoV-2/B.1, SARS-CoV-2/BA.5, MERS-CoV, HCoV-229E, HCoV-NL63, HCoV-OC43 and mock controls. Arrows represent trajectories based on RNA velocity and signal strength is represented by the length of the arrows.

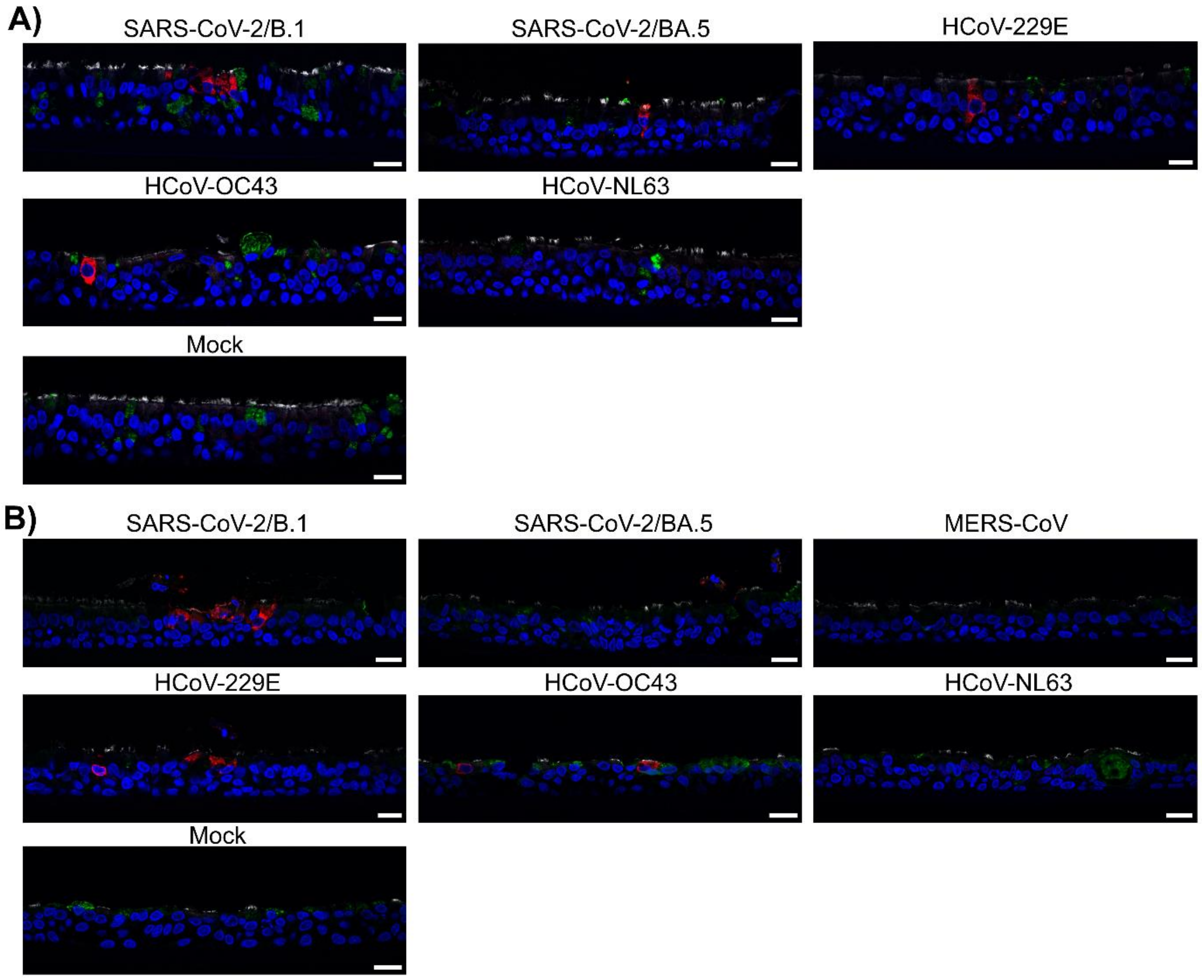


**Supplementary Figure 5: Human coronaviruses show preferential tropism for ciliated epithelial cells.** Representative immunohistochemistry stain of BCi **(A)** and SABCi **(B)** fixed at 4 dpi following virus infection. Sections were immunostained to identify infected cells (viral nucleocapsid, red), ciliated cells (α-Tubulin, white), and or goblet cells (Mucin 5AC [MUC5AC], green). Sections were counter stained with DAPI (blue). Images are representative of two independent experiments. Scale bar = 20 μm.

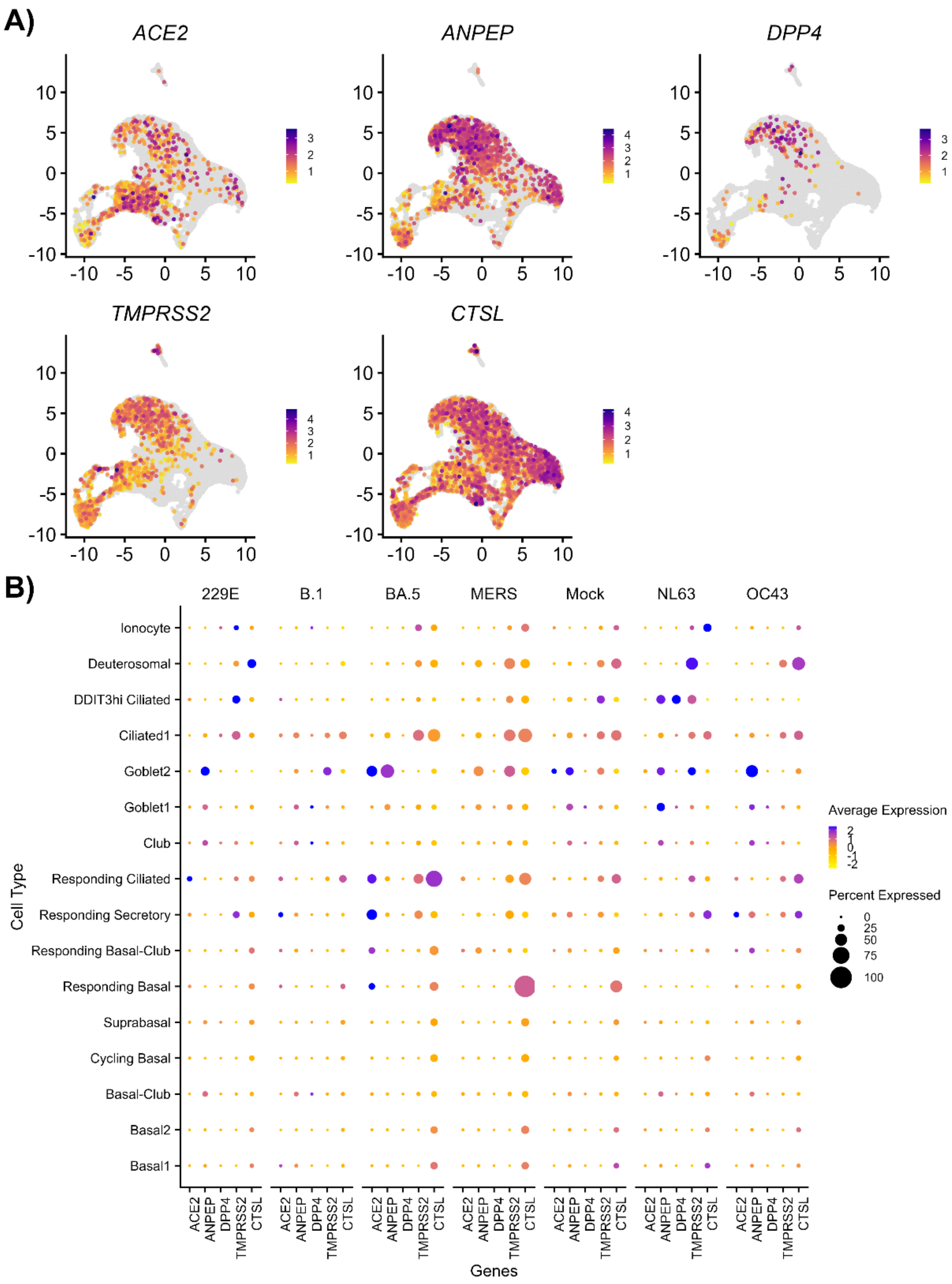


**Supplementary Figure 6: Expression of entry factors in human nasal epithelial cells. (A)** UMAP representation of cell profiles from each experimental condition, coloured by normalised expression of ACE2, DPP4, ANPEP, TMPRSS2 and CTSL. Colour scale indicates the mean of normalized expression of marker genes in each gene. **(B)** Violin plot of HCoV receptor and entry factors genes for distinct cell types in the mock condition.

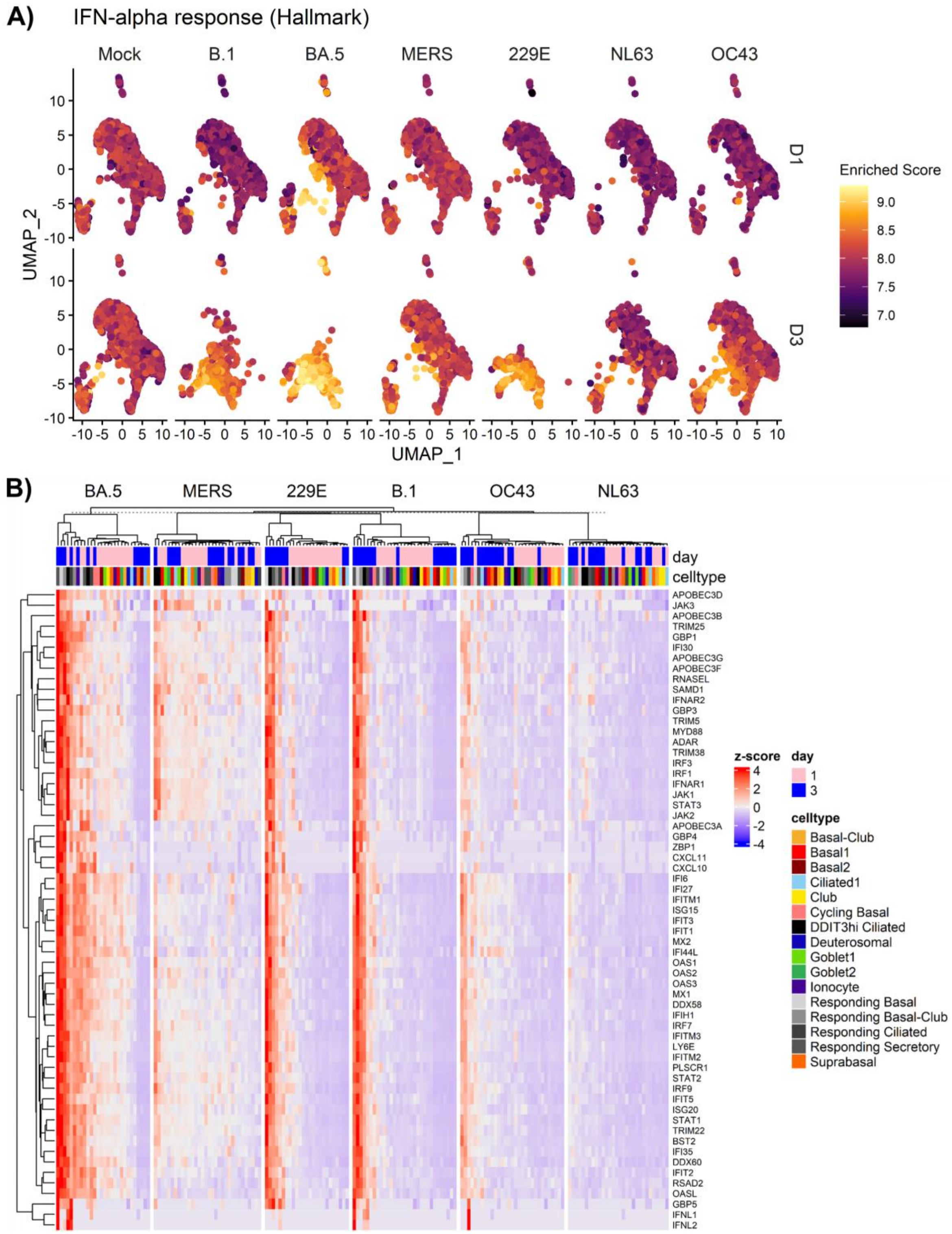


**Supplementary Figure 7: Interferon responses to human coronaviruses in human nasal epithelial cells. (A)** UMAP representation of cell profiles from each experimental condition, coloured by enriched IFN-alpha response hallmark score. **(B)** Z-score of selected genes associated with interferon signalling in each condition, cell type and day post infection.

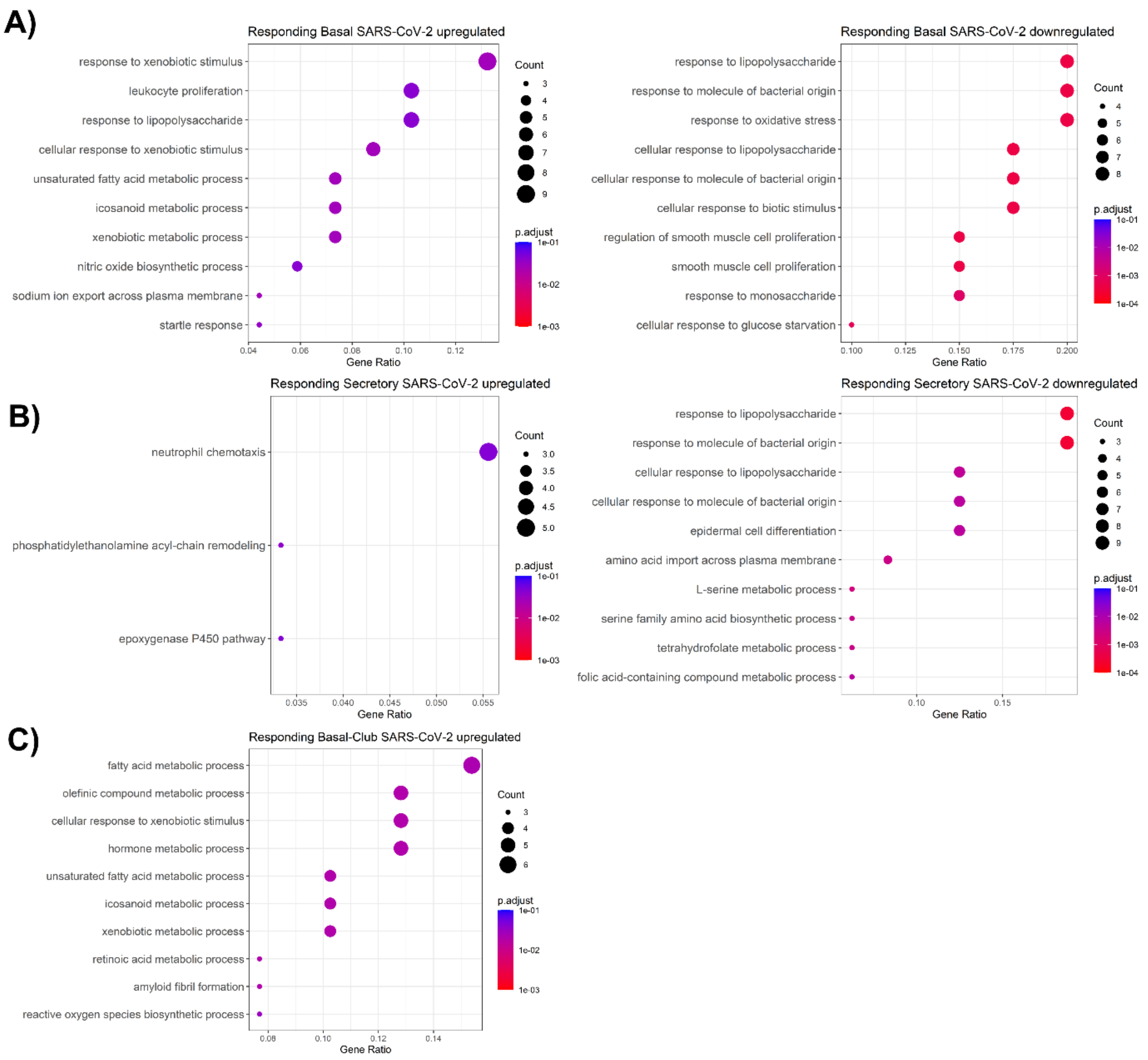


**Supplementary Figure 8: Pathway analysis of differentially expressed genes in responding cell populations between SARS-CoV-2 and HCoV-229E exposed cells.** Top 10 Gene Ontology Biological Pathways enriched responding basal **(A)**, responding basal-club **(B)** and responding secretory cells **(C)**. Colour indicates adjusted p-value and size of dot indicates number of genes in each pathway.

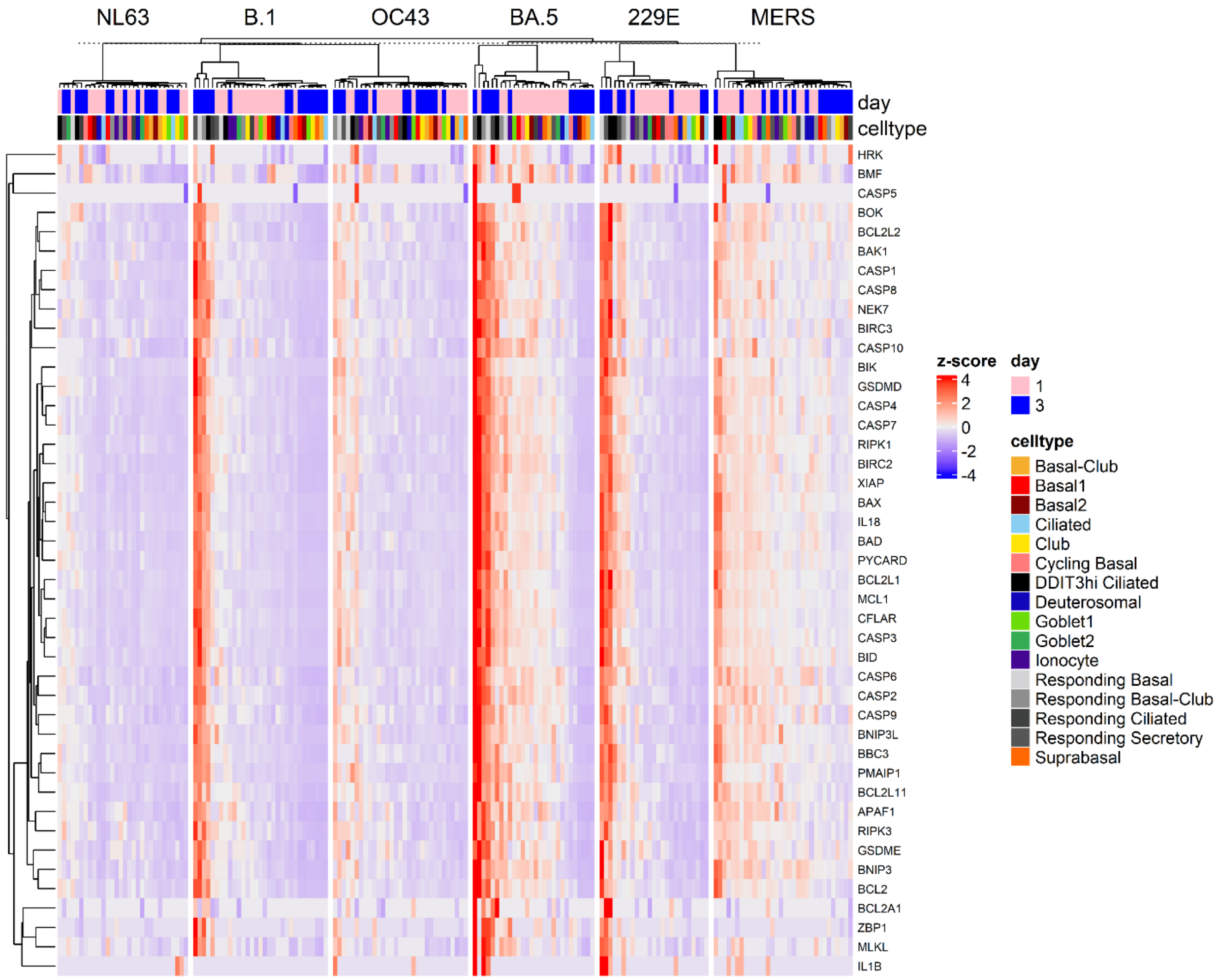


**Supplementary Figure 9: Induction of genes associated with cell death following human coronavirus infection in nasal epithelial cells.** Z-score of selected genes associated with cell death in each condition, cell type and day post infection.

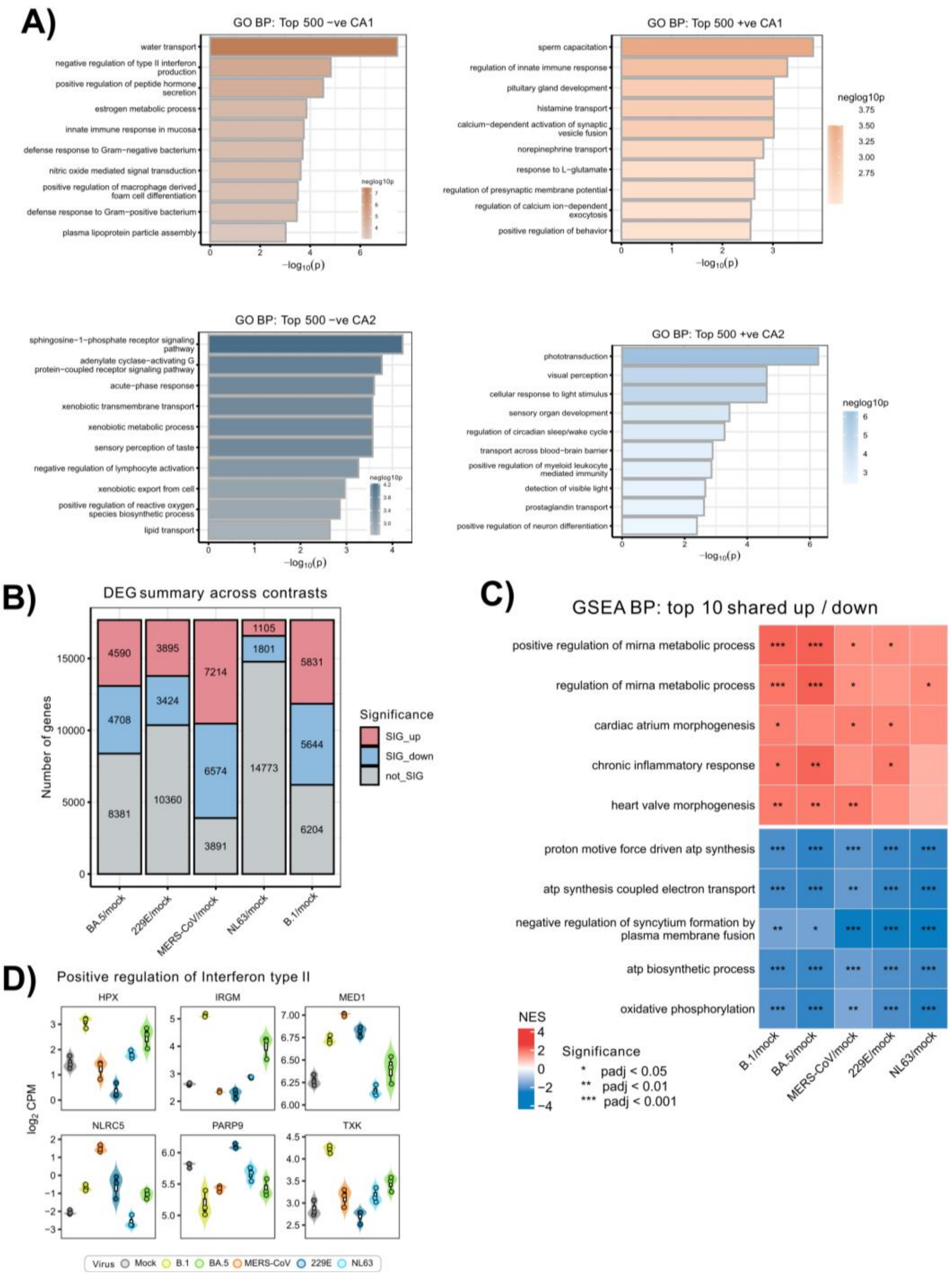


**Supplementary Figure 10: Analysis of gene expression in edAT2 cells following human coronavirus infection.** **(A)** Correspondence analysis (COA) of all conditions with top GO BP enrichment of the top 500 positive and negative genes in CA1 and CA2. **(B)** The number of upregulated and downregulated differentially expressed genes (DEGs) across all differential expression tests (contrasts). **(C)** GSEA of top shared Biological Processes gene terms across each of the 5 DGE analyses (each viral infection against mock) * P adj. < 0.05, ** P adj. < 0.01, P adj. < 0.001 **(D)** Gene expression signatures of positive regulation of type II interferon.
